# YORU: social behavior detection based on user-defined animal appearance using deep learning

**DOI:** 10.1101/2024.11.12.623320

**Authors:** Hayato M Yamanouchi, Ryosuke F Takeuchi, Naoya Chiba, Koichi Hashimoto, Takashi Shimizu, Ryoya Tanaka, Azusa Kamikouchi

## Abstract

The creation of tools using deep learning methodologies for animal behavior analysis has revolutionized neuroethology. They allow researchers to analyze animal behaviors and reveal causal relationships between specific neural circuits and behaviors. However, the application of such annotation/manipulation systems to social behaviors, in which multiple individuals interact dynamically, remains challenging. Here, we applied an object detection algorithm to classify animal social behaviors. Our system, packaged as “YORU” (Your Optimal Recognition Utility), classifies animal behaviors, including social behaviors, based on the shape of the animal as a “behavior object”. It successfully classified several types of social behaviors ranging from vertebrates to insects. We also integrated a closed-loop control system for operating optogenetic devices into the YORU package. YORU enables real-time delivery of photostimulation feedback to specific individuals during specific behaviors, even when multiple individuals are close together. We hope that the YORU system will accelerate the understanding of the neural basis of social behaviors.

## Introduction

Social behaviors such as courtship, aggression, and group formation are important for improving survival rate and reproductive efficiency in a wide range of animals, including invertebrates and vertebrates ^1–3^. To accelerate our understanding of the neural basis of these behaviors, it is necessary to capture information on the location and type of social interactions that occur between individuals ^4^. Recent advances in machine learning have led to the establishment of various tools for animal behavior detection, represented by markerless body part tracking ^5–7^. These emerging tools enable real-time behavior analysis, allowing researchers to manipulate neural activity precisely when the animal exhibits the behavior of interest ^8–10^. Such a closed-loop approach promises to clarify the causal relationship between the neural activity and behaviors, and indeed has yielded significant success in uncovering neural basis for single individual behavior ^7,11,12^. However, detection of the social interaction between multiple individuals with low latency remains challenging ^4,6,7^. This difficulty arises from the complexity of defining social behavior based on the coordinates of each individual’s body parts, preventing the effective use of a closed-loop approach ^6,7,13^. Therefore, novel methods are required to enable the detection of social interaction and to perform real-time behavior analysis for neural intervention.

Social interactions are often accompanied by individuals striking distinctive postures to interact with others, such as wing extension by courting male fruit flies and mounting on females by male mice attempting copulation ^14–16^. In this study, we propose the detection of social behaviors of multiple individuals based on their appearances by defining each as a “behavior object”. To this end, we focused on an object detection algorithm “YOLOv5”, a high-speed object detection algorithm based on a convolutional neural network (CNN) ^17,18^. The algorithm achieves fast object recognition by framing detection as a single regression problem, using a unified architecture that processes the entire image in one forward pass ^17,18^. The YOLO-based detection is robust to variations in object orientation, size and background noise ^19^. YOLOv5 is therefore a promising algorithm to detect various animals’ social behaviors without relying on the body-part coordinates, making it well-suited for adapting behavior analysis and a closed-loop system to investigate social interactions.

Here, we established a behavior detection system called “YORU” (Your Optimal Recognition Utility), based on the YOLOv5 algorithm ^18^. First, to verify the proof of concept, we tested whether YORU could be used to detect social behaviors of various animals ranging from vertebrates to insects. As an application of our behavior analysis, we compared the detection readouts with neural activity imaging in mice to interpret large-scale brain activity. Secondly, we evaluated the inference speed of the detection and feedback latency in YORU’s real-time analysis. Finally, as a practical example of the YORU’s closed-loop system, we experimented with optogenetic neural manipulation focused on courtship behavior in *Drosophila*. In the presence of multiple flies, we successfully manipulated the neural activity of individuals in an individual selective manner by optogenetic stimulation of the fly exhibiting the behavior of interest.

## Results

### YORU is a framework for animal behavior recognition

Here we introduce “YORU” (Your Optimal Recognition Utility), an animal behavior detection system with a graphical user interface (GUI). In the YORU system, animal behaviors, either performed by single animals or multiple animals interacting with each other, are classified as a “behavior object” based on their shapes with YOLOv5 ^18^ (Fig. 1a). YORU is an open-source Python software composed of four packages: “Training”, “Evaluation”, “Video Analysis” and “Real-time Process” (Fig. 1b, Figs. 1a-e). The YORU system is designed to work both offline (video analysis) and online (real-time analysis) to detect animal behavior. During video analysis, YORU allows us to quantify animal behavior according to user-defined shapes of social behavior. For real-time analysis, YORU analyzes animal behavior in real-time, which can be used to output trigger signals accordingly to control external devices, such as LEDs for optogenetic control as set by the user.

**Fig. 1:**
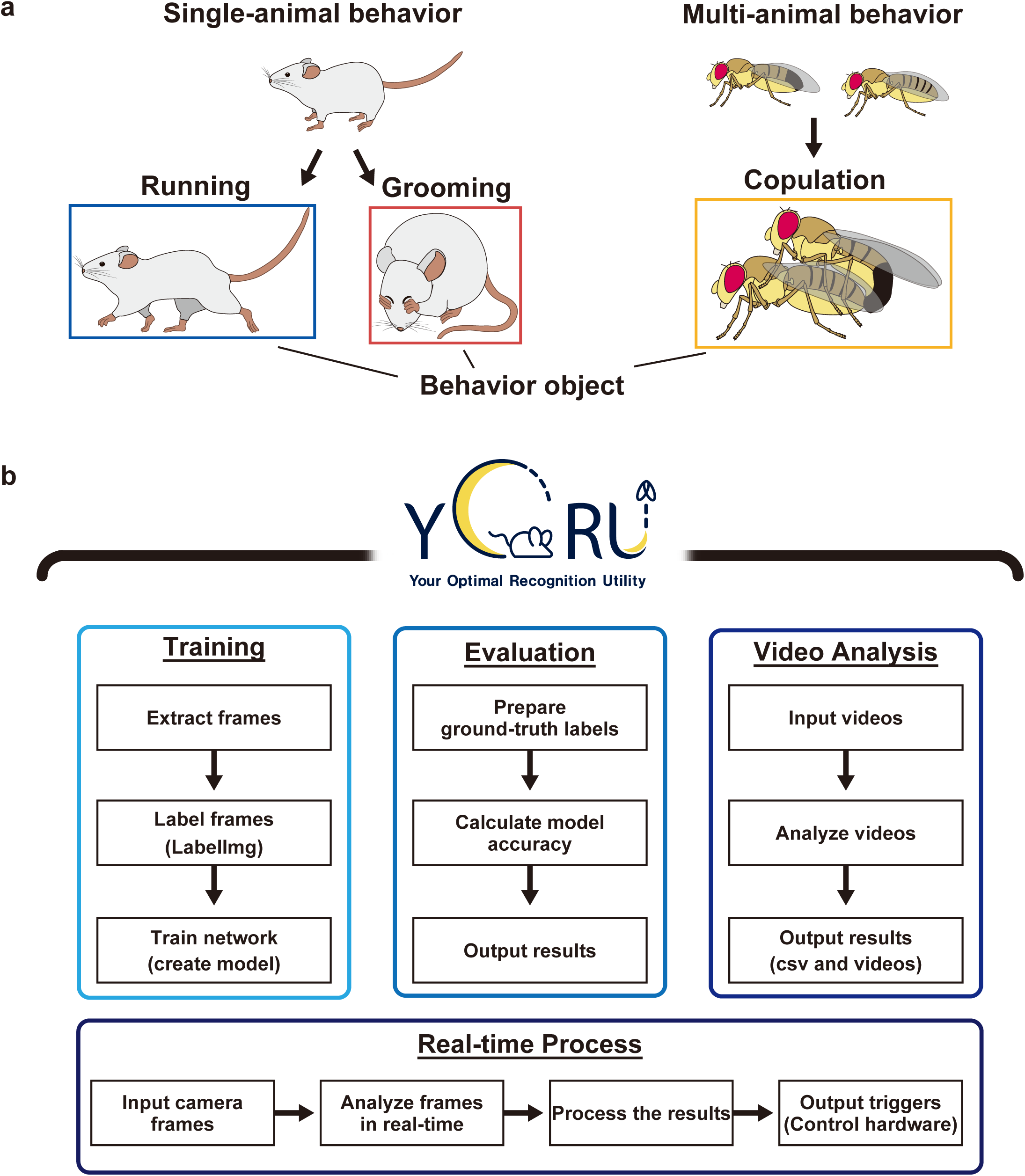
YORU detects animal behaviors as a behavior object. **a**, Illustrations of behavior objects. YORU can adapt to single-animal (Left) and multi-animal (Right) behaviors, including social behaviors. Animal behaviors (Top) can be classified as behavior objects (Bottom). **b**, Diagram of YORU. YORU includes four packages: Training, Evaluation, Video Analysis, and Real-time Process.

We set three constraints to the workflow of YORU: ease of use for experimenters, low system latency, and high customizability. To make YORU user-friendly, we designed the system to allow easy quantification of animal behaviors without requiring programming knowledge. To achieve low system latency, special efforts were focused on behavior detection and feedback when operating as a closed-loop system. Here, immediate behavior detection is achieved by YOLOv5-based object detection algorithm, which processes generating region proposals and classifying subjects simultaneously, resulting in faster detection ^17,18^. Immediate feedback is achieved by the parallel processing design of YORU real-time process ^20^: image acquisition, object recognition, and hardware (Arduino, DAQs, etc.) manipulation are not processed serially but simultaneously. To achieve high customizability, i.e., adapting YORU system to various experimental systems easily, YORU implements a trigger output for hardware manipulation. The experimenter can customize YORU’s parameters with no or minimal user programming. These features enable experimenters to easily use high-performance closed-loop neural feedback.

### Detection of social behaviors using object detection algorithm

In previous studies, YOLO-based object detection successfully classified some social behaviors of *Drosophila*, such as offensive behaviors and copulation ^21,22^. In this study, we extended this idea and analyzed a variety of animal behaviors with YORU. The following social behaviors were tested to validate the performance: (1) Fruit flies; wing extension behavior of a male toward a female during courtship ^14^ (Fig. 2a), (2) Ants; mouth-to-mouth food transfer behavior among workers (trophallaxis) ^23,24^ (Fig. 2b), and (3) Zebrafish; orientation behavior toward another individual behind a partition ^25,26^ (Fig. 2c). We extracted 2000 images from multiple videos and manually labeled their behavior objects with the following definitions.

1. Fruit flies: “wing_extension”; a fly extending one of its wings (Fig. 2a). When the flies were not labeled as “wing_extension”, they were labeled as “fly”.
2. Ants: “trophallaxis”; heads of two ants engaging in a food exchange, with their mouth parts in contact with each other. “no”; the situation of no food exchange although the heads of the two ants were close together (Fig. 2b). No label was given when neither of these two behavior types were detected.
3. Zebrafish: “orientation”; two zebrafish exhibiting orientation behavior as defined in a previous study ^25,26^, “no_orientation”; a zebrafish exhibiting no orientation behavior (Fig. 2c).

**Fig. 2:**
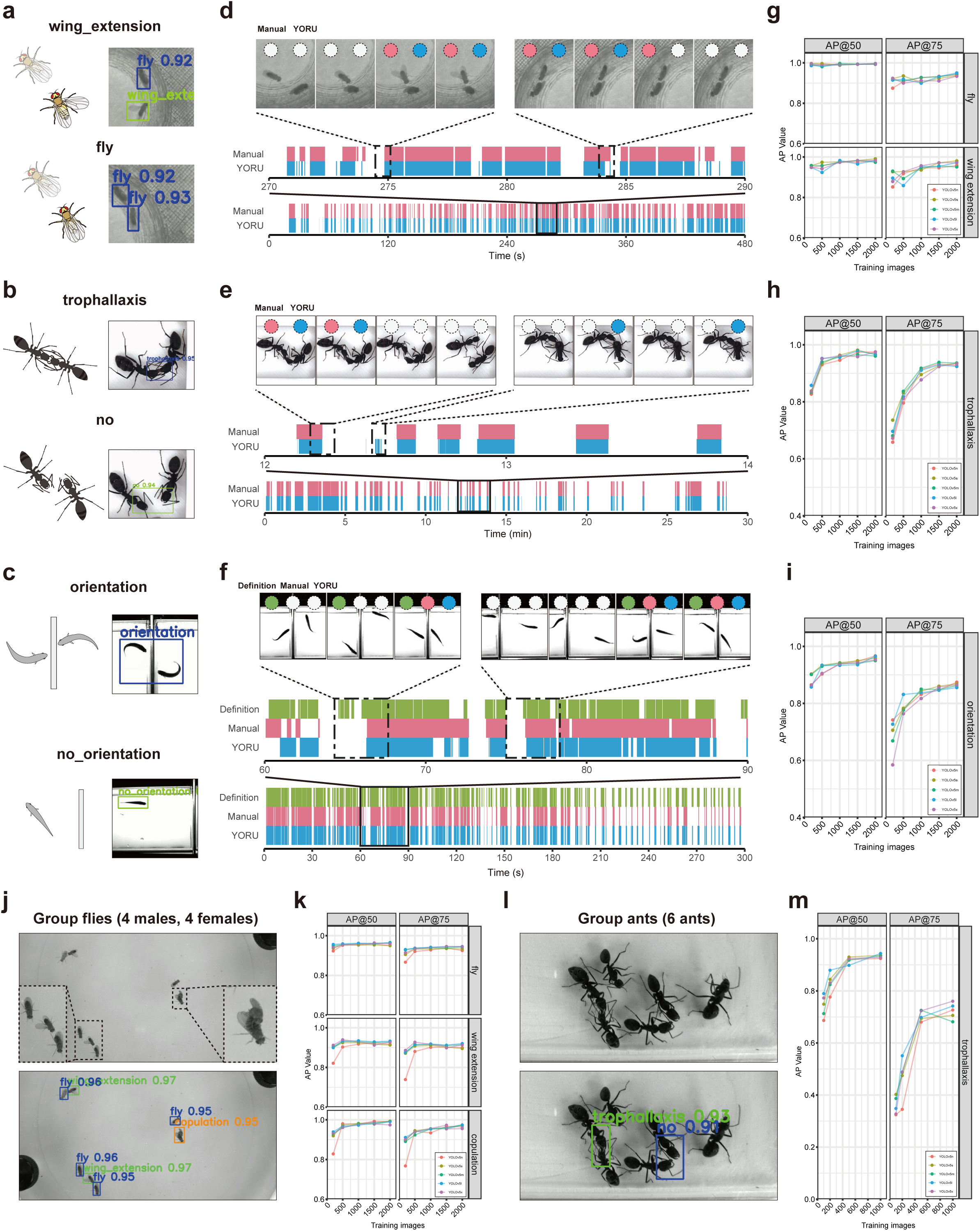
Detection of animal behaviors by YORU. **a**, “Fly - wing extension” dataset. Two behavior classes were defined: “wing_extension”; a fly extending one of its wings, and “fly”; a fly that does not belong to the “wing extension” class. **b**, “Ant - trophallaxis” dataset. Two behavior classes were defined: “trophallaxis”; two ants showing nutrition exchange, and “no”; the situation of no nutrition exchange but the heads of the two ants are close together. **c**, “Zebrafish - orientation” dataset. Two behavior classes were defined: “orientation”; two zebrafish showing orientation behavior, and “no_orientation”; zebrafish showing no orientation behavior. **d-f**, Ethogram of the fly wing extension (**d**), ant trophallaxis (**e**), and zebrafish orientation (**f**) behaviors. Top panels show the example view of each behavior. The circles show detections by manual analysis (pink), YORU analysis (blue), or analysis using a previous tracking method (green). The horizontal axes in the ethogram represent the observation period. The colored area of the ethogram shows the occurrence of the behavior detected with YORU analysis (blue), manual analysis using BORIS (pink), and the previous tracking method (green). **g-i**, The AP values of “Fly - wing extension” (**g**), “Ant - trophallaxis” (**h**), and “Zebrafish - orientation” (**i**) models. The values at IOU=50% (AP@50) (Left) and IOU=75% (AP@75) (Right) are shown. The horizontal and vertical axes show the number of images used for training and the AP values, respectively. Each colored line shows the AP values of different pre-trained models (Also in **k**, **m**). **j**, “Fly - group courtship” datasets. Three behavior classes were defined: “copulation”; a pair of flies showing copulation, “wing_extension”; a fly extending one of its wings, and “fly”; a fly that does not belong to the “copulation” and “wing extension” classes. **k**, The AP values for “Fly - group courtship” models. **l**, “Ant - group trophallaxis” dataset. Two behavior classes were defined: “trophallaxis”; two ants showing nutrition exchange, and “no_trophallaxis”; the heads of two ants are close together without exchanging nutrition.

We created models and compared their detection accuracies with human annotations using multiple videos that were not used for model creation. The Accuracy score for the model’s detection of fruit flies, ants, and zebrafish behaviors were 93.3%, 98.3% and 90.5%, respectively, when compared to human manual annotations (Fig. 2d-f, Supplementary Table 1). We also compared zebrafish orientation analyses between the previous method based on body part tracking ^25^ and human annotations, yielding an Accuracy score of 81.2% (Supplementary Table 1, Fig. 2f). YORU can thus potentially detect zebrafish orientation quicker and with higher accuracy than the previous tracking methods. These results suggest that YORU’s detection can detect social behaviors with a similar to human annotations.

In general, two major factors to consider for the practical application of deep learning-based analyses are the amount of training data (the number of labels) and the selection of based networks ^5,27^. To find optimal conditions in our dataset, we evaluated the accuracy of models with different numbers of training images (200, 500, 1000, 1500, and 2000) and YOLOv5 networks (YOLOv5n, YOLOv5s, YOLOv5m, YOLOv5l, and YOLOv5x). Two metrics were used in this evaluation: “Precision”, the ratio of correctly predicted detections to all predicted detections, and “Recall”, the ratio of correctly predicted detections to all ground-truths. All models detected the target social behavior with Precision greater than 85% and Recall greater than 90%, even when only 200 images were used to create models (Supplementary Table 2). The Precision and Recall scores increased with the number of training images, finally exceeding 90% and 95%, respectively, when 1,000 or more images were used for training (Supplementary Table 2). We then evaluated typical object detection model indices, intersection over union (IOU) and average precisions (AP) (Fig. 2g-i, Figs. 2a-c) ^28^. These two indexes can be used to compare the accuracy of the model between conditions; the IOU shows the difference in location information between the ground truth and detected bounding boxes while the AP reflects the accuracy of the behavior classes, which includes Precision and Recall values ^28^. For each behavior, neither the number of images used for training nor the YOLOv5 models tested affected IOU values (Figs. 2a-c). On the other hand, AP values increased with the number of training images, in which the same number of images gave similar AP values across YOLOv5 based models (Fig. 2g-i). These evaluations suggest the primary factor influencing accuracy is the amount of training data, at least in these cases.

Next, we increased the number of individuals to test the performance of YOLOv5 models in multi-individual conditions. In the first condition, a group of flies was analyzed to detect three objects: “wing extension”, “copulation”, and other behaviors labeled as “fly” (Fig. 2j, k). In the second condition, a group of ants was used to detect two objects: “trophallaxis” and “no_trophallaxis” (Fig. 2l, m). In the group of flies, Precision and Recall scores were over 95% even when using only 200 images for training; the scores increased with the number of training images (Supplementary Table 2). In the group of ants, Precision and Recall scores exceeded 90% with 500 or more training images (Supplementary Table 2). These findings suggest that YORU’s detection of social behaviors is also applicable to multiple individuals (more than two individuals) with high accuracy (Fig. 2 j-m, Figs. 3a, b, Supplementary Table 2), highlighting its usefulness for analyzing various types of social behaviors. The detection accuracy does not change significantly with different YOLOv5 pre-trained models and instead depends strongly on the number of training images.

**Fig. 3:**
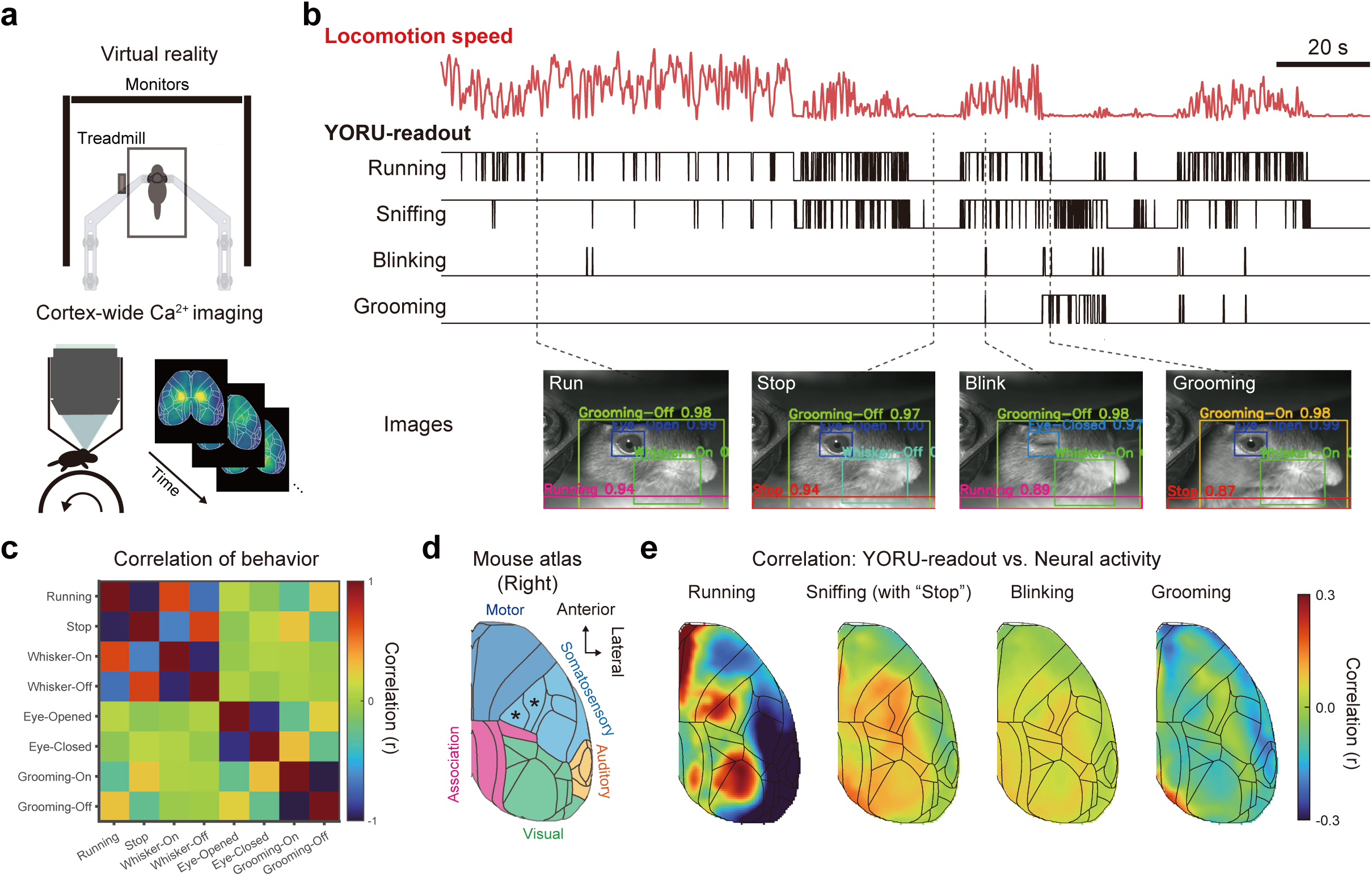
YORU uncovers the relationship between behavioral readouts and neural activity interpretation. **a**, The setup for virtual reality (VR) and cortex-wide imaging in mice. **b**, Time-series of locomotion speed (red, calculated from rotary encoder signals of VR) and YORU readout (black). Representative images are shown at the bottom. **c**, Correlation matrix between each behavior. Correlation indices are derived from the YORU readout time series data. **d**, Top view of the Allen Common Coordinate atlas of the dorsal cortex. Five rough divisions are auditory areas (yellow), association areas (magenta), somatosensory areas (cyan), and motor areas (blue). Black asterisks show the somatosensory areas representing forelimb and hindlimb information. **e**, Pseudo-colormap of Spearman’s correlation coefficient (YORU-readout vs. Neural activity of each pixel).

### The relationship between behavioral readouts and neural activity interpretation

One of the main questions in neurophysiology is the interpretation of which sensations and animal behaviors can explain observed neural activity. One promising methodology to address this question is to combine time series analyses of multiple behavioral types *via* video analysis with neural activity measurements. To test the performance of YORU for such tasks, we recorded dorsal cortex-wide neural activity with wide-field calcium imaging from mice running in a virtual reality (VR) system which provides visual feedback coupled to their locomotion ^29^ (Fig. 3a). A mouse on a treadmill in VR typically exhibits multiple behaviors, such as running, grooming, and eye blinking (Fig. 3b). First, to validate the behavior classification performance of YORU, we estimated the time series of eight behavior classes from video analysis: “Running”, “Stop”, “Whisker-On”, “Whisker-Off”, “Eye-Open”, “Eye-Closed”, “Grooming-On”, and “Grooming-Off”. Precision and Recall scores for the model’s detection of these behaviors were 91.8% and 92.7%, respectively, validating this model to detect mouse behaviors as accurately as human manual annotations (Supplementary Table 3). For all classes detected by this model, the average IOUs and AP@50 (i.e., the AP value at IOU=50) were above 0.60 and 0.55, respectively (Fig. 4, Supplementary Table 3). Previous studies have reported that rodents actively move their whiskers to seek and identify objects or avoid obstacles in front of them during locomotion ^30–33^. In line with these reports, the time series data of active whisker movement (corresponding to “Whisker-On” label) during “Running” periods were positively correlated while “Whisker-Off” and “Running” correlated negatively (Fig. 3b, c). In addition, the “Running” and “Stop” epochs in the YORU readout are negatively correlated, indicating that mutually exclusive behaviors are labeled exclusively (Fig. 3c). These results further confirmed that YORU’s approach to behavior labeling can efficiently detect typical behaviors of mice.

**Fig. 4:**
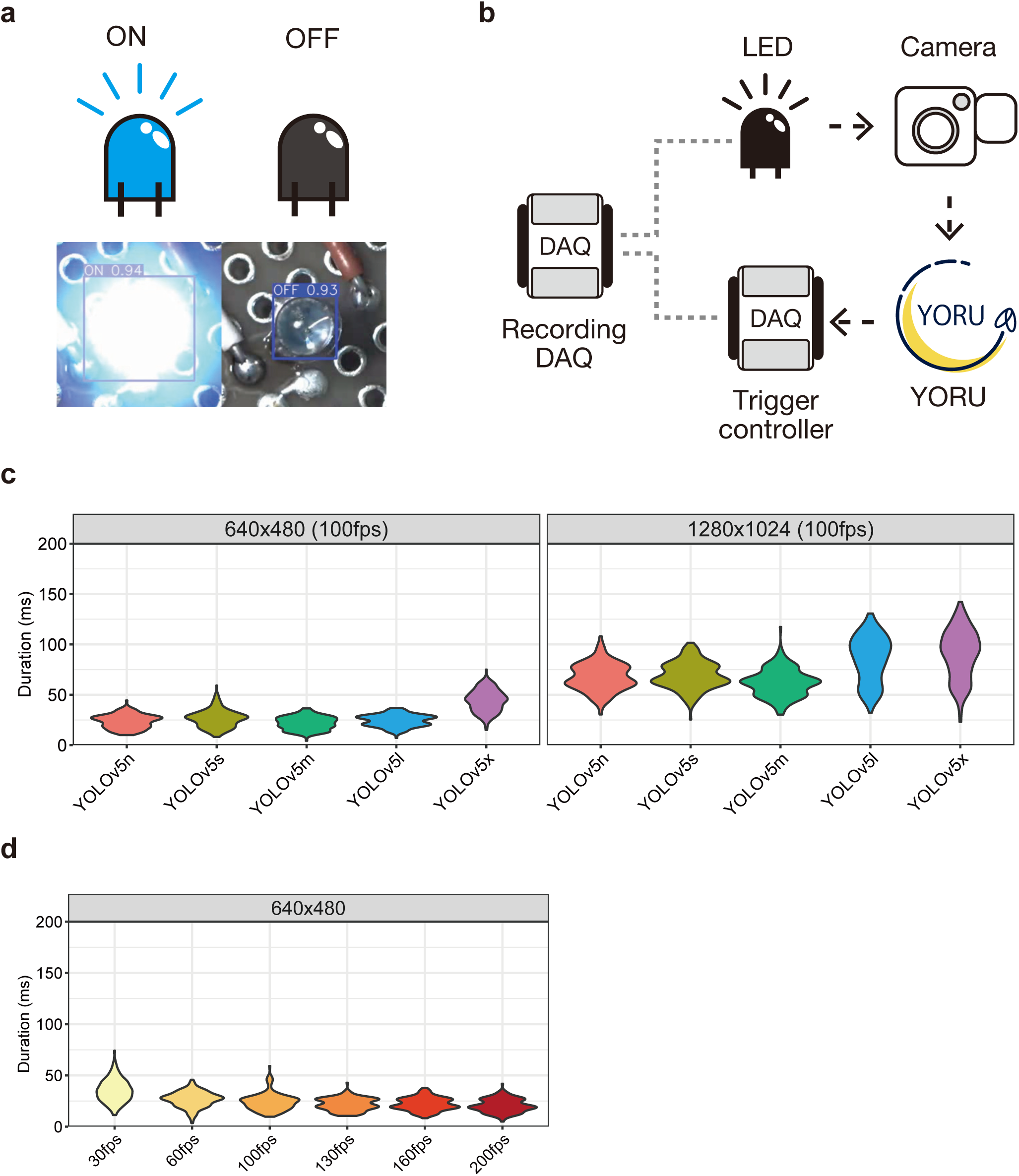
Validation of YORU’ s operation speed. **a**, “LED lighting” dataset. Two state classes were defined: “ON” and “OFF”, indicating that LED light was turned on and off, respectively. **b**, Schematic of system latency measurements. The camera captures the LED light, YORU detects a frame, and the trigger-controller DAQ outputs the TTL voltage based on detection by YORU. The recording DAQ logged the TTL pulse from the trigger controller DAQ and the LED voltage. **c**, The system latency of “LED lighting - Small” (Left) and “LED lighting - Large” (Right) models. The system latency of each model was calculated using camera images with resolutions of 640×480 pixels and 1240×1024 pixels, respectively. **d**, The system latency at different camera frame rates. The “LED lighting - Small” model was used. **c**,**d**, Violin plot represents the probability density of individual data points within the range of possible values.

Next, we investigated which brain regions in the cortex correlate with YORU’s readout. The “Running” epoch was highly correlated with the neural activity of medial motor areas, somatosensory areas representing forelimb and hindlimb information, visual areas, and posterior association region (retrosplenial cortex) (Fig. 3d). Several whisker movements associated with sniffing during “Stop” period and “Blinking” behavior were also coupled to distinct macroscopic activity patterns of widespread regions in somatosensory and visual areas. As expected, grooming behavior correlated specifically with neural activity of forelimb somatosensory and motor areas (Fig. 3d, e). These results demonstrate the applicability of its utility and potential for precise and quantitative interpretation of neural activity with various animal behaviors.

### Inference speed and system latency of YORU

A closed-loop system that relies on live feedback of animal behaviors requires a low-latency solution for behavior detection and feedback outputs ^12^. To assess YORU’s potential for application in closed-loop systems for social behavior analysis, we measured the total time required from frame acquisition to behavior estimation using a simple light detection task (Fig. 4a). Possible major factors that could affect the speed of YORU’s behavior detection are (i) the network structure, (ii) image size, and (iii) computing hardware, especially the graphics processing unit (GPU). To demonstrate their impact on analyzing each frame, we measured the single-frame inference latency with (i) five YOLOv5 architectures (YOLOv5n, YOLOv5s, YOLOv5m, YOLOv5l, and YOLOv5x), (ii) two input image sizes (640×480 or 1280×1024 pixels, images were resized before feeding to the neural network), and (iii) a variety of NVIDIA GPUs in Windows PCs (Supplementary Table 3). Inference speed was calculated by comparing the time before (t1) and after (t2) the frame was analyzed by a model (Figs. 5a). We found that the smallest network (YOLOv5n) was the fastest and the largest network (YOLOv5x) had the largest inference latency (Figs. 5b, Supplementary Table3). In addition, the inference speed was faster with a smaller image size and more powerful NVIDIA GPUs (Supplementary Table 3). For example, with NVIDIA RTX 4080 GPU, we achieved inference speed as low as ∼ 5.0 ms per frame (∼ 200 fps) using YOLOv5s networks (Figs. 5b). These results suggest that the network architecture, input image size, and GPU all influence YORU’s inference speed.

**Fig. 5:**
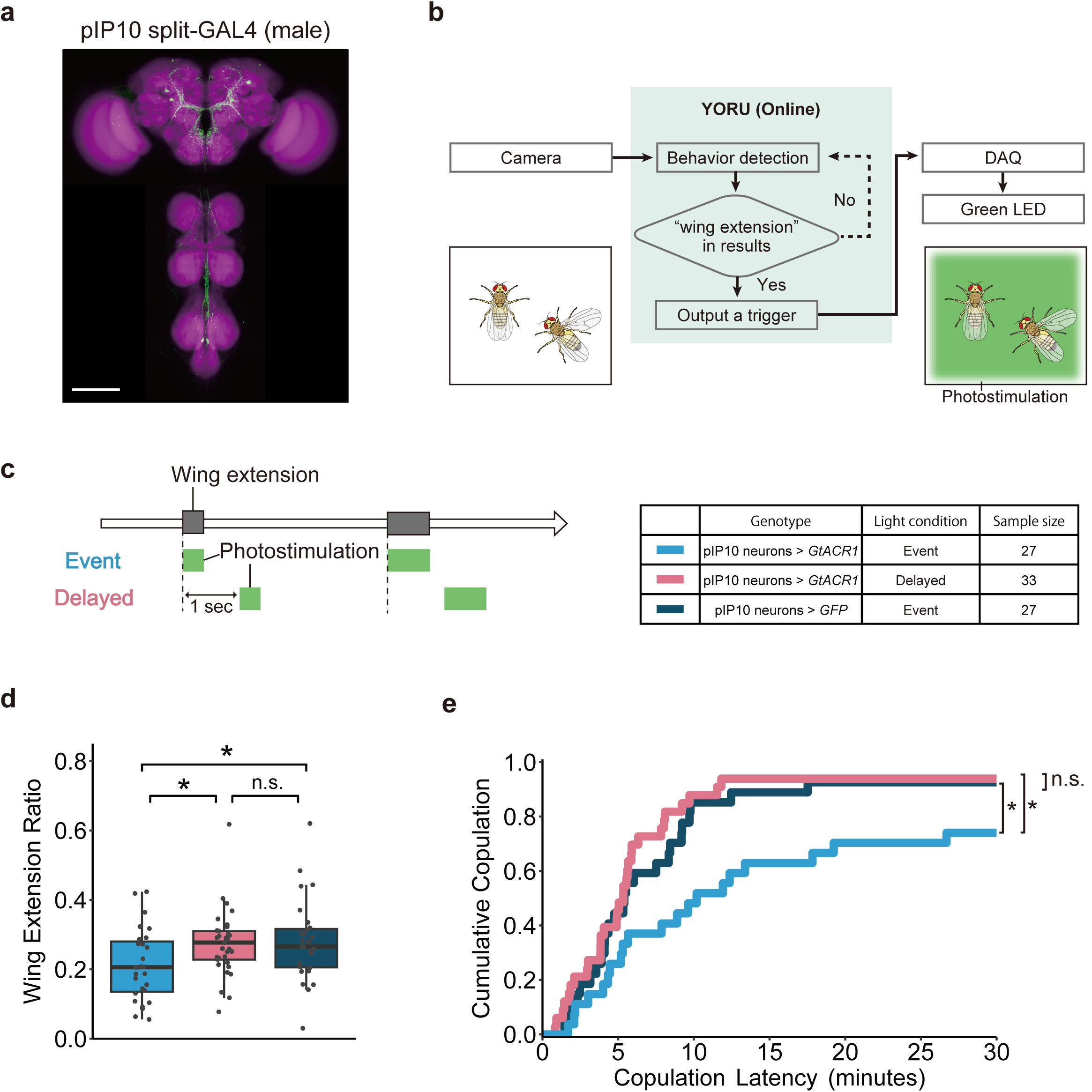
Neural manipulation in response to male wing extension using YORU. **a**, Microscopy image of pIP10 split-GAL4 expression pattern in a representative male brain and ventral nerve cord. Scale bar, 100 μm. Signals of the GFP marker (green) and counter-labeling with the nc82 antibody (magenta) are shown. **b**, Schematic of the YORU’ s closed-loop conditions. YORU analyzes camera frames. Then, if YORU detects “wing extension”, it sends signals to the trigger controller (DAQ) and operates the LED light. **c**, Schematic of light conditions. As the experimental photostimulation condition, YORU introduces green photostimulation to the entire chamber when it detects a fly showing wing extension (Event). As a photostimulation control, we used an event-triggered light with a 1-second delay that illuminates the entire chamber (Delayed). pIP10 specific split-*GAL4>UAS-GtACR1* (pIP10 neurons > *GtACR1*) males were used for these two groups. In addition, pIP10 split-*GAL4>20XUAS-IVS-mCD8::GFP* (pIP10 neurons > *GFP*) males were used as a genetic control. The following color code was used: experimental group (blue), photostimulation control group (pink), and genetic control group (dark blue) (Also in **d**, **e**). **d**, Ratio of time spent displaying wing extension before copulation. The aligned rank transform one-way analysis of variance (ART one-way ANOVA) test corrected with the Benjamini-Hochberg method was used for statistical analysis. Boxplots display the medians (horizontal white line in each box) with 25th and 75th percentiles and whiskers denote 1.5x the interquartile range. Each point indicates individual data. **e**, Cumulative copulation rate of pIP10 split-*GAL4>GtACR1* males. Pairwise comparisons using Log-Rank test corrected with the Benjamini-Hochberg method were used for statistical analysis. **d**,**e**, Not significant (n.s.), p > 0.05; *, p < 0.05.

The design of closed-loop systems adapted to neuroscience experiments requires the simultaneous control of several processes, such as camera capture and hardware manipulation ^34^. In addition, several other factors such as camera type and frame rate, trigger destination type, and PC memory can affect system latency. Therefore, we tested the performance of YORU’s closed-loop system from end to end, including the entire process from the camera capturing an image of the LED to the PC detecting whether the LED was lit or not and sending a trigger signal to a data acquisition (DAQ) system upon detecting the “ON” state (Fig. 4a, b). The delay between the timing of the LED turning on and the trigger signal output, both detected by measuring their voltages using DAQ, was as low as 30 ms per event (Fig. 4c). This suggests that the end-to-end system latency of the YORU system is around 30 ms in this setup, which is sufficiently low in most cases to provide real-time feedback in response to animal behavior. Next, we assessed the factors that affected the end-to-end system latency, such as networks, input image size, camera frame rate, and system hardware. The effect of the network differences was almost negligible except YOLOv5l and v5x, which showed larger latency than others (Fig. 4c). On the other hand, the input image size severely affected the latency; the average latency of the smaller image (640 x 480 pixels) was ∼30 ms while that of the larger image (1280 × 1024 pixels) was ∼75 ms (Fig. 4c). According to the camera fps latency results, even if the camera frames were acquired as fast as the model’s inference speed (∼200 fps), the system delay would not be significant, due to YORU’s multi-processing system (Fig. 4d). In a multi-processing system, the size and processing speed of the random access memory (RAM) affect the processing speed of the system due to the shared memory. Interestingly, in YORU’s real-time process, the size of the RAM (between 16 GB and 32 GB) had less effect on the system latencies, suggesting that 16GB RAM size is minimally sufficient to operate YORU’s real-time process (Figs. 6a). The effect on system latency due to hardware differences (cameras and trigger devices) was small (Figs. 6b, c). These results suggest that inference speed and the input image size are the primary factors affecting end-to-end system latency. In addition, YORU’s system processing speed was very fast compared to previously reported tracking-based systems ^7^.

**Fig. 6:**
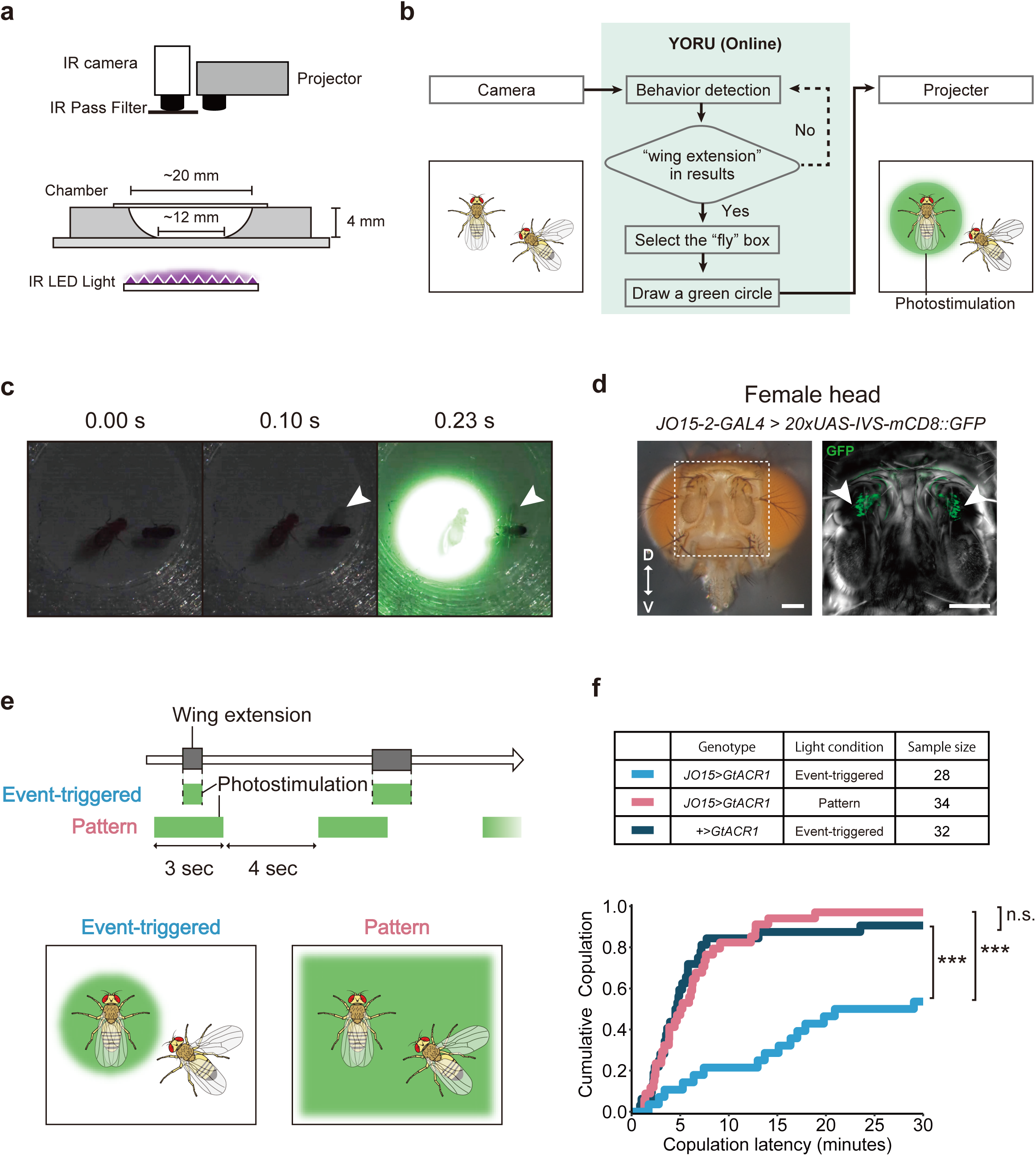
Individual-specific neural manipulation by YORU. **a**, Experimental setup of individual-focused photostimulation. An infrared (IR) camera is used for observing flies. IR LED light is used for IR camera recording. IR pass filter allows only IR light to be captured by the IR camera, preventing visible light noise. The projector is used for introducing individual-focused photostimulation. **b**, Schematic of the closed-loop system for individual-focused photostimulation assay. YORU analyzes camera frames. If YORU detects “wing extension”, it draws a green circle on the “fly” bounding box and sends the image to the projector. The projector introduces green-circled photostimulation to the fly. **c**, A representative situation during individual-focused photostimulation. White arrowheads show the fly displaying wing extension. **d**, *JO15-2-GAL4* expression in female antenna. GFP markers driven by *JO15-2-GAL4* (*JO15-2-GAL4>20XUAS-IVS-mCD8::GFP*) are detected in Johnstons Organs. White arrowheads show JO neurons. D and V indicate the dorsal and ventral sides, respectively. Scale bar, 100μm. **e**, Schematic of light conditions. In the experimental photostimulation condition, when YORU detects a fly showing wing extension, YORU introduces individual-focused photostimulation to the other fly (Event-triggered). In the photostimulation control, we used pattern light (3s On, 4s Off) to illuminate the entire chamber irrespective of the wing extension event (Pattern). **f**, Cumulative copulation rate of *JO15-2>GtACR1* females. We used *JO15-2-GAL4>UAS-GtACR1* (*JO15-2>GtACR1*) females for the experimental group. As a genetic control, we used *+>UAS-GtACR1* (*+>GtACR1*) females. The following color code was used: experimental group (blue), photostimulation control group (pink), and genetic control group (dark blue). Pairwise comparisons using Log-Rank test corrected with the Benjamini-Hochberg method used for statistical analysis. Not significant (n.s.), p > 0.05; ***, p < 0.001.

### YORU application for real-time optogenetic system

Next, to assess the practical applicability of the YORU closed-loop system, we applied it to event-triggered optogenetic manipulation. During courtship, the male fruit fly extends his wing to serenade the female with a unique sound known as the courtship song. Upon hearing it, female flies gradually increase their receptivity to copulation ^35^. We hypothesized that if male wing extension behavior is inhibited when a male attempts to extend his wing, copulation rates would be reduced. Using a split-GAL4 strain that specifically drives gene expression in pIP10 neurons, which are descending neurons regulating courtship song production ^36,37^, we expressed the green light-gated anion channel GtACR1 ^38^ in male pIP10 neurons (Fig. 5a). We then paired individual mutant males with a wild-type female in a chamber and allowed YORU to detect the single-wing extension of a fly. In this system, when YORU detects an object labeled as “wing extension,” YORU introduces green photostimulation to the entire chamber (Fig. 5b, c). As a control for photo-stimulation, we used an event-triggered light with a 1-second delay that illuminated the entire chamber (Fig. 5c, ‘Delayed’ group). Male flies expressing GFP in pIP10 neurons were also used as a genetic control group. We then analyzed the wing extension ratio during courtship and the copulation rate during the 30-min observation period (Fig. 5d, e). In the experimental group, males decreased the amount of wing extension during courtship (Fig. 5d), validating the optogenetic inhibition of pIP10-induced behavior. In line with this, the cumulative copulation rate was significantly lower in the experimental group than in the control groups (Fig. 5e). These results confirm the importance of male pIP10 neurons for inducing the wing extension behaviors observed in the previous reports ^36,37,39^, which subsequently leads to male copulation success. They also validate YORU’s performance in operating real-time manipulation of neural activity in response to the detection of a specific behavior.

### YORU application for individual-focused photo-stimulation

Finally, we applied YORU for individual-selective neural manipulation in response to social behavior between multiple individuals. We included a projector in YORU’s closed-loop system to control the light pattern for optogenetic stimulation (Fig. 6a-c). When YORU detects the target behavior, it sends location information to focus the projector light. We utilized the fly courtship assay to test the usability of the system by suppressing female hearing only when the male sings a courtship song, as exhibited by his wing extension. We hypothesized that disrupting hearing in females during male courtship song production would suppress female mating receptivity, leading to a reduced copulation rate. Using the *JO15-2-Gal4* strain that selectively labels auditory sensory neurons, we expressed GtACR1 ^38^ in female auditory sensory neurons (i.e., JO-A and JO-B neurons) (Fig. 6d, Fig. 7). We then paired these females each with a wild-type male in a chamber and utilized the online capabilities of YORU. In this system, when YORU detects an object labeled “single-wing extension,” the YORU-operated projector illuminates the object labeled as “fly”, which typically is the female courted by the male (Fig. 6b, c, e). As a control for individual-focused photo-stimulation, we used pattern light (3s On, 4s Off) illuminating the entire chamber (Fig. 6e). In the experimental group (*JO15-2>GtACR1* female with event-triggered light condition), the copulation rate was significantly lower than in the control groups (Fig. 6e, f). This result confirms the importance of auditory sensory neurons for females to detect the male’s courtship song to enhance copulation receptivity. Again, YORU was able to optogenetically manipulate neural activity using individual-focused illumination, even when multiple individuals were moving in a chamber at the same time. Those experiments validated the usefulness of the YORU system, which can manipulate various devices, such as a projector as well as a DAQ and Arduino, by trigger output. These proof-of-principle experiments demonstrate the usefulness of YORU for the online detection of social behaviors and manipulation of individual-focused neural activity through optogenetics.

## Discussion

Here, we presented YORU, an animal behavior detection system using a YOLOv5-based object detection algorithm. YORU allowed the detection of social behaviors, as well as single-animal behaviors. Furthermore, by introducing real-time analysis, YORU can operate a closed-loop system with low latencies and high user scalability. We also demonstrated the practical applicability of real-time neuronal manipulation using the fly courtship behavior. In particular, we created an individual-focused illumination system to manipulate the neural activity of selected individuals in response to a specific behavior. The YORU’s closed-loop system is thus a powerful approach for social behavior research. On the user side, YORU can be used entirely through its GUI without any programming. Just as body parts tracking-based analysis tools such as DeepLabCut ^5^ have revolutionized neuroscience research, YORU will meet the needs of many biologists and stimulate the generation of novel, testable hypotheses.

In tracking-based behavioral analyses, behavior is typically defined based on the location of body parts. In addition, clustering analysis (e.g., t-SNE, PCA, UMAP) of body part location coordinates can be used to classify behaviors and detect unknown behavior patterns ^5^. However, it is challenging for tracking-based analysis to classify already known behaviors with high accuracy in real-time. For example, tracking-based analysis for recognizing behaviors such as wing extension in *Drosophila* requires recognition of wings and body axes and definition of behaviors by their angles, and accordingly, it would fail to capture behaviors if body parts are not correctly tracked even partially. In addition, to capture the behavior of multiple individuals, we need to know which body part belongs to which animal: Despite various algorithms being proposed to overcome these challenges, capturing the behavior of multiple individuals in real time is still difficult ^6,7,13^. YORU’s object detection algorithm can sufficiently compensate for the shortage of body part tracking-based behavior quantification even in multiple animal conditions ^6,7^. YORU’s processing time is fast enough for real-time processing, even at camera resolutions sufficient for animal behavior quantification.

In behavioral neuroscience, optogenetics serves as a powerful approach to cell-type- and spatiotemporal-specific control of neural activity, especially for investigating the causal relationship between neural circuits and behavior ^8,40^. With state-of-the-art genetic tools, it is now possible to express channelrhodopsin only in specific neurons to perform neural activity intervention ^8,38^. In addition, the development of computer science has made it possible to create closed-loop experimental setups and manipulate neural activity during behavior, allowing us to explore the causal relationship between neural circuits and behavior in more detail ^34,41,42^. In the research of the neural bases of social behaviors, optogenetic manipulation of only specific individuals, even in the presence of multiple individuals, can be a powerful approach ^43^. However, the difficulty of capturing the behavior of multiple individuals simultaneously in real-time has made it extremely difficult to conduct online behavior analysis in the presence of multiple individuals. The YORU system allows us to create a closed-loop system that can analyze animal behavior and operate photostimulation for optogenetics in real-time. Furthermore, the YORU system allows us to conduct individual-focused optogenetics experiments with widely available equipment: projectors, cameras, and personal computers.

Due to YORU’s concept of detecting behaviors by their “snapshot” appearances, it has limitations in detecting behaviors that cannot be defined within a single frame and instead require time-series data for definitions ^21^, such as foraging or mating attempts. To capture these behaviors, it is necessary to adapt object detection algorithms to these time-series behaviors, or to perform additional analysis using other tools such as DeepLabCut. Another limitation lies in the hardware operation. Although the system delay in our condition was ∼30 msec, an additional delay in trigger processing depending on the external hardware should be considered. In our experiments of individual-focused phtostimulation, there was an additional delay on the projector side before the patterned image signal was sent and projected. This time lag may cause fast-moving individuals to move out from the area of the light stimulus. Possible solutions for such cases include incorporating predictive algorithms, or using a high-speed projector such as a low latency gaming projector ^44,45^. Overcoming these two limitations would further widen the potential options for studying dynamic animal social behaviors.

Various analysis tools, driven by advances in deep learning, have contributed to biology research. YORU is a user-friendly system that allows all analyses to be performed *via* a GUI, making it easy for anyone to perform analysis both in offline and online modes. The object detection paradigm has the potential not only for animal behavior quantification, but also for capturing various biological phenomena. In particular, it has been incorporated as a tracking method for humans and fish, and has been applied to the behavioral classification of animals and plants (e.g., *Drosophila* mating behaviors and the stomatal opening and closing of *Arabidopsis*) ^21,22,46,47^. The YORU system can contribute to making the use of object recognition algorithms more widespread in biology and to significantly reduce the labor of biologists.

## Materials and Methods

### Development of YORU system

YORU is written in Python. The GUI of YORU is developed using DearPyGui library, which enabled GUI development in a fast, interactive manner, and plotting of acquisition data in real-time. To use various image acquisition devices such as webcams or other high-performance machine vision cameras, we used OpenCV API for image acquisition. All processing and data streams were performed in a multiprocessing manner with the Python library “multiprocessing”. To label animal behaviors, YORU uses the open-source annotation software “LabelImg” (https://github.com/HumanSignal/labelImg). To detect “behavior objects”, YORU uses YOLOv5 packages (https://github.com/ultralytics/yolov5). The code for YORU is available at (https://github.com/Kamikouchi-lab/YORU) as open-source software. In this study, we used ChatGPT (https://chatgpt.com), developed by OpenAI, as a tool to assist in various programming tasks. ChatGPT was used for code generation, debugging, refactoring suggestions, and answering technical questions. While ChatGPT was used to assist in specific tasks, all generated output was reviewed and validated by the researchers to ensure accuracy and reliability.

### Datasets

To evaluate the performance of YORU, we prepared different datasets collected under various conditions. The datasets include “Fly - wing extension”, “Ant - trophallaxis”, “Fly - group courtship”, “Ant - group trophallaxis”, “Zebrafish - orientation”, “LED lighting - Small”, and “LED lighting - Large”. The “Zebrafish - orientation” dataset was created using zebrafish videos from a previous study ^26^, while the other datasets were generated from videos obtained in this study. Each dataset consisted of images (frames manually extracted from videos) and “behavioral object” labels. Each image was then manually labeled using LabelImg. Supplementary Table 5 shows the detailed condition of each dataset.

#### Flies - wing extension

This dataset includes a pair of wild-type fruit flies consisting of a male and a female. Fruit flies (*Drosophila melanogaster*, Canton-S strain) were raised on standard yeast-based media on a 12 h light/12 h dark (12 h L/D) cycle. Both sexes of flies were collected within 8 h after eclosion to ensure their virgin status. They were maintained at 25°C under a 12 h L/D cycle and transferred to new tubes every 2 to 4 days, except on the day of the experiment. Male flies were kept singly in a plastic tube (1.5 mL, Eppendorf) containing ∼200 μL fly food, while females were kept in groups of 10 to 30. Experiments were conducted using males and females 4 to 8 days post-eclosion, with each individual used only once. The video recordings were performed between Zeitgeber Time (ZT) = 1-11 at 25°C and 40-60% relative humidity.

Courtship behavior was monitored in a round courtship chamber with a sloped wall (20 mm top diameter, 12 mm bottom diameter, 4 mm height, and 6 mm radius fillet) made of transparent polylactic acid filament by 3D printer (Sermoon D1, Creality 3D Technology Co., Ltd.). The chamber was enclosed with a slide glass and a white acrylic plate as a lid and bottom, respectively. The chamber was illuminated from the bottom by an infrared LED light (ISL-150×150-II94-BT, 940 nm, CCS INC.) to enable recordings in dark conditions. Male and female flies were gently introduced into chambers by aspiration without anesthesia. Videos were captured from the top, with a monochrome CMOS camera (DMK33UX273, The Imaging Source Asia Co., Ltd.) equipped with a 25 mm focal length lens (TC2514-3MP, KenkoTokina Corporation) and a light-absorbing and infrared transmitting filter (IR-82, FUJIFILM), at a resolution of 640 x 480 pixels and 30 fps for 30 min for each pair using IC Capture (The Imaging Source Asia Co., Ltd.). In this dataset, we labeled two behavior object classes, “fly” and “wing_extension”, without a female or male identification. The “wing_extension” was labeled on the fly when it extended one of its wings. The “fly” indicates a fly not showing wing extension.

#### Flies - group courtship

This dataset includes a group of wild-type fruit flies consisting of four males and four females. We prepared these flies under the same condition described in the “Fly - wing Extension” dataset. Behaviors were monitored in a fly bowl chamber with a sloped wall (60 mm in diameter, 3.5 mm in depth) ^48^, illuminated by a visible LED light to facilitate recordings. Four male and four female flies were gently introduced into the chamber by aspiration without anesthesia. Videos were captured from the top as described in the “Fly - wing Extension” dataset. In this dataset, we labeled three behavior object classes: “fly”, “wing_extension”, and “copulation” without a female or male identification. The definitions of “wing_extension” and “fly” were based on those in the “Fly - wing extension” dataset. The “copulation” was defined by genital coupling between a male and a female.

#### Ants - trophallaxis

This dataset includes two worker ants. Workers of *Camponotus japonicu*s were collected in October 2023 at the Higashiyama Campus of Nagoya University (35°09’15.3” N, 136°58’15.8” E). Ant species were identified with the *Encyclopedia of Japanese Ant* ^49^. After collection, they were kept individually in Falcon tubes with moistened paper for 24 h at 22°C with 40-60% relative humidity under dark conditions. The video recordings were conducted at 22°C and 40-60% relative humidity.

The trophallaxis behavior was monitored in a custom-made chamber (25 mm width, 15 mm length, 5 mm depth) made of transparent polylactic acid filament by 3D printer (Sermoon D1, Creality 3D Technology Co., Ltd.). The chamber was enclosed with a glass ceiling. Two worker ants were used for a single video recording: one individual was fed 1 M sucrose solution as much as desired immediately before observation, while the other was not fed anything. These two workers were transferred to a custom-made chamber. Videos were captured from the top, with a color CMOS camera (DFK33UP1300, The Imaging Source Asia Co., Ltd.) equipped with a 50 mm focal length lens (MVL50M23, Thorlabs, Inc.), at a resolution of 1280 x 960 pixels and 30 fps for 30 min for each pair using IC Capture (The Imaging Source Asia Co., Ltd.). In this dataset, we labeled two behavior object classes: “trophallaxis” and “no”. The “trophallaxis” class was defined as a situation when the heads of two individuals were in proximity and the palps were in contact with each other. The “no” class was defined when the heads of the two individuals were in proximity, but the palps were not in contact with each other.

#### Ant - group trophallaxis

This dataset includes a group of ants consisting of six worker ants. Among six workers, three individuals were fed 1 M sucrose solution as much as desired immediately before observation, while the other three were not fed anything. These workers were then transferred to the polystyrene chamber (87mm width, 57mm length,19mm depth) with a glass ceiling, and video recording was started. Videos were recorded from the top with a monochrome CMOS camera (FLIR GS3-U3-15S5; Edmund optics) equipped with a zoom lens (M0814-MP2, CBC Optics Co., Ltd.). Video recordings and behavior object definitions were performed in the same manner as for the “Ant - trophallaxis” dataset.

#### Zebrafish - orientation

This dataset includes a pair of wild-type zebrafish. Adult zebrafish, aged four to six months, with the Oregon AB genetic background were used without sex identification. Fish were maintained in a 14/10 h L/D cycle at 28.5°C. The experiments were conducted as described in previous studies ^25,26^. Fish were isolated in individual tanks the day before the experiments, with white paper placed between the tanks to prevent visual contact. The experiments were conducted at ZT = 0-14.

The orientation behavior was monitored in two custom-made acrylic tanks (90 mm length, 180 mm width, and 60 mm height). These tanks were separated by a divider made from a polymer dispersed liquid crystal (PDLC) film (Sunice Film) attached to 2 mm thick acrylic sheets. Fish were placed individually into water tanks with a depth of 57 mm for 20 min. Following this acclimation period, the fish were recorded for 5 min at 30 fps with the opaque divider condition (invisible condition). The divider was then made transparent to allow the fish to see each other, and the recording continued for an additional 5 min (visible condition). Videos were captured from below, with a monochrome CMOS camera (FLIR FL3-U3-13E4, Edmund optics) at a resolution of 1280 × 1024 pixels and 30 fps. The divider condition (opaque or transparent) was controlled with a DAQ interface (USB-6008; National Instruments Co.) and custom-made software written in LabVIEW (National Instruments Co.). In this dataset, we labeled two behavior object classes: “orientation” and “no_orientation”. The “orientation” class was defined as a situation when two zebrafish showed orientation behavior as defined in a previous study ^25^. The “no_orientation” class was defined when two zebrafish showed no orientation behavior (Fig. 2f).

#### LED – ON or OFF

This dataset includes a blue LED (OSB5YU3Z74A, OptoSupply). We created “LED lighting - Small” and “LED lighting - Large” datasets; “LED lighting - Small” consists of 640×480 pixel videos, and “LED lighting - Large” consists of 1280×1024 pixel videos. Videos were captured with a color CMOS camera (DFK33UP1300, The Imaging Source Asia Co., Ltd) equipped with a 50 mm focal length lens (MVL50M23, Thorlabs, Inc.). In each dataset, we created models based on each YOLOv5 model (YOLOv5n, YOLOv5s, YOLOv5m, YOLOv5l, and YOLOv5x). In these models, there were two classes: “ON” and “OFF”, indicating that LED turned on or off, respectively (Fig. 4a).

### Model creation for YORU-based analysis

We created YORU models using the “Training” package. YORU randomly splits the datasets containing the images and labels into two datasets: 80% for training and 20% for validation. The models were trained with the training and validation datasets for 300 epochs. If the training loss was low enough, training was finished before 300 epochs. For comparison with human manual annotations, we created YORU models based on a YOLOv5s pre-trained model, in which 2000 images were used. Human manual annotations were conducted using BORIS ^50^. For model evaluation using “Evaluation” package, we extracted the specified number of images from each dataset and created each model based on YOLOv5 pre-trained models.

### Comparison with YORU and human manual annotations in animal videos

To obtain human manual annotation data, we analyzed the behaviors of flies (10 videos and ∼40 min in total), ants (3 videos and ∼90 min in total), and zebrafish (2 videos and ∼10 min in total), following the behavior definitions (Supplementary table 5). Four parameters, Accuracy, Precision, Recall, and F1 score, were used to compare the performances between the human manual annotation and YORU. To obtain values for these parameters, each annotation was first classified as follows:

・True Positive (TP): Correct annotation.

・False Positive (FP): Incorrect annotation, such as annotating a non-existing object or a misplaced annotation.

・False Negative (FN): Undetected ground-truth.

・True Negative (TN): Correct no-annotation.

Subsequently, Accuracy, Precision, Recall, and F1 score were calculated as follows:

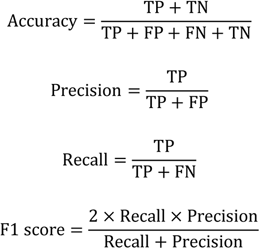

### Evaluation of YORU models

We evaluated the models using YORU’s “Evaluation” package. Several hundred images, which were not used in the model creation, were manually labeled with ground-truth bounding boxes using LabelImg. YORU then predicted bounding boxes on the same images using the models. By comparing the ground-truth and predicted boxes, we calculated the Precision and Recall of the models. In addition, intersection over union (IOU) and average precisions (AP), two of the typical object detection model indexes, were used ^28^. IOU represents how close the ground-truth box and the predicted box are, based on Jaccard Index that evaluates the overlap between two bounding boxes, and AP represents the accuracy of the behavior classes, reflecting the “Precision” and “Recall” values ^28^. Since the AP is affected by the IOU threshold, we used two different threshold values to calculate the APs: “AP@50” with a threshold value of IOU = 50% and “AP@75” with a threshold value of IOU = 75%. The AP ranges from 0 to 1, with a value of 1 indicating perfect consistency with the ground-truth labels. As with the PASCAL Visual Object Classes Challenge ^51^, model evaluation in YORU adopts the AP using the all-point interpolation method by the Riemann integral to compare the models.

### Mice – running in virtual reality and recording neural activity

All mice procedures followed institutional and national guidelines and were approved by the Animal Care and Use Committee of Nagoya University. All efforts were made to reduce the number of animals used and minimize the suffering and pain of the animals. The “Mouse - treadmill” dataset consists of 3 videos of single mouse behaviors in the treadmill environment. Referring to a previous study ^29^, the videos were collected. C57BL/6J mice were purchased from Nihon SLC. These animals were maintained in a temperature-controlled room (24°C) under a 12 h light/dark cycle with ad libitum access to food and water. Due to the body size needed for adaptation in head-fixed VR, mice aged 11−28 weeks were used in the VR experiments. The VR environment used in this study was referred to in the previous study ^29^. Briefly, mice were head-fixed and free to run on a one-dimensional treadmill. We recorded mice behavior with a monochrome CMOS camera (DMX33UX174, The Imaging Source) at 30 fps and 640 × 480 pixels resolution.

For the “Mouse - treadmill” dataset, we labeled 2171 frames with eight behavior object classes: “Running”, “Stop”, “Whisker-On”, “Whisker-Off”, “Eye-Open”, “Eye-Closed”, “Grooming-On”, and “Grooming-Off” (Fig. 3b). “Eye-Open” and “Eye-Closed” indicate a mouse that opens or closes the eye. “Grooming-On” was defined as the situation where the forelegs of the mouse are touching its head. “Grooming-Off” was defined as all situations that do not satisfy the criteria of “Grooming-On”. “Running” and “Stop” indicate a mouse that runs or stops. “Running” was defined as the situation where the roller was rotating, and “Stop” was defined as the situation where the roller was stationary. “Whisker-On” was defined as the situation where the whisker was oriented in the anterior direction, and “Whisker-Off” was defined as all situations that did not satisfy the criteria of “Whisker-On”. Labels were randomly split into two datasets: creating a model dataset (1975 images) and a test dataset (196 images). Using creating a model dataset, we generated a model based on the YOLOv5s pre-trained model.

Surgery for the head plate implantation and virus injection were performed as previously described ^29^. Briefly, Retro-orbital virus injection, The AAV-PHP.eB-hSyn-jGCaMP7f was injected at 100 µL into the retro-orbital sinus of mice using a 30-gauge needle. Head plate implantation was performed 14 days after the virus injection. Skin and membrane tissue on the skull were carefully removed, and the surface was covered with clear dental cement (204610402CL, Sun Medical). A custom-made metal head-plate was implanted onto the skull for stable head fixation. During surgery, the eyes were covered with ofloxacin ointment (0.3%) to prevent dry eye and unexpected injuries, and body temperature was maintained with a heating pad. All procedures of surgery were performed under deep anesthesia with a mixture of medetomidine hydrochloride (0.75 mg/kg; Nihon Zenyaku), midazolam (4 mg/kg; Sandoz), and butorphanol tartrate (5 mg/kg; Meiji Seika). After surgery, mice were injected with atipamezole hydrochloride solution (0.75 mg/kg; Meiji Seika) for rapid recovery from the effect of medetomidine hydrochloride.

The recording of cortex-wide neural activity was performed as previously described ^29^. Briefly, we used a tandem lens design (a pair of Plan Apo 1×, WD=61.5 mm, Leica). To filter the calcium independent artifacts ^52^, an alternating blue (M470L4, Thorlabs, Inc.) and violet (M405L3, Thorlabs, Inc.) excitation LEDs were used as a light source for excitation, combined with band-pass filters (86-352, Edmund; FBH400-40, Thorlabs, Inc., respectively). Fluorescent emission was passed through a dichroic mirror (FF495-Di03) and filters (FEL0500 and FESH0650, Thorlabs, Inc.) to a scientific CMOS (sCMOS) camera (ORCA-Fusion, Hamamatsu Photonics). The timing of the excitation light was controlled by a global exposure timing signal from the camera, processed with an FPGA-based logic circuit (Analog discovery 2, Digilent), equipped with binary-counter IC (TC4520BP, Toshiba).

### Speed benchmarking – inference speed

We measured the inference speed of the YOLOv5 detection using a custom Python code of the YORU’s detection function in the “Real-time Process” package. The list of computers used for this analysis is shown in Supplementary Table 6. In the custom Python code, the times before and after the YOLO detection step (*t*1 and *t*2, respectively) were logged by the “perf_counter()” function of the “Time” module. Then, we calculated the inference speed of one frame detection as *t*2 − *t*1. For the analyses on the model size and frame size dependency, we used the 640 × 480 pixels videos for “LED lightning - Small” models and 1280 × 1024 pixels videos for “LED lightning - Large” models. 50000 frames (60 fps, ∼14 min, ∼420 events) were used to calculate the inference speed.

### Speed benchmarking – real-time system

To estimate the latency of the YORU’s “Real-time Process” package, we measured the end-to-end latency of the LED light detection task by running YORU on a Windows desktop PC (CPU, Core i7-13700KF 16core; GPU, NVIDIA RTX 4080; RAM, 32GB or 16GB DDR5). The CMOS camera (DFK 33UP1300, The Imaging Source Asia Co., Ltd or ELP-USBFHD08S-MFV, Autocastle) equipped with a 50 mm focal length lens (MVL50M23, Thorlabs, Inc.) captured an LED and streamed frames. The frames were then processed to detect whether the LED state was on or off using the “LED lighting - Small” or “LED lighting - Large” model. When YORU detected the ON state, it sent a signal to a trigger controller, which then emitted a transistor-transistor logic (TTL) voltage pulse. As the trigger controller, DAQ (USB-6008, National Instruments Co.) or a microcontroller (Arduino Uno, Arduino CC) was used. A recording DAQ (USB-6212, National Instruments Co.) logged the TTL voltage from the trigger controller (Fig. 4b). The voltage data were then processed using a 50 Hz low-pass filter (“butterworth_filter()” function of “scipy” package) to block the high-frequency noises. The delay between the timing of LED voltage and that of the trigger TTL was used to estimate the full-system latency, which includes overhead from hardware communication and other software layers.

### Optogenetic assays – fly preparation

*D. melanogaster* were raised as described above. Canton-S was used as a wild-type strain. *UAS-GtACR1.d.EYFP (attP2)* ^38^ (RRID: BDSC_32194) and split-GAL4 strain that labels pIP10 neurons specifically (*w; VT040556-p65.AD; VT040347-GAL4.DBD*) ^36^ (RRID: BDSC_87691) were obtained from the Bloomington Drosophila Stock Center. *JO15-2-GAL4* ^53^ was a kind gift from Dr. D. F. Eberl (University of Iowa).

For optogenetic assays, pIP10 neurons specific *split-GAL4>GtACR1* male or *JO15-2>GtACR1* female flies were paired with wild-type adult males or females, respectively, as mating partners. Flies used for the behavior assay were collected within 8 h after eclosion to ensure their virgin status. Wild-type male, transgenic male and female flies were kept singly in a plastic tube (1.5 mL, Eppendorf) containing ∼200 μL fly food. Wild-type females were kept in groups of 10 to 30. They were transferred to new tubes every 2 to 3 days, but not on the day of the experiment. Males and females, 5 to 8 days after eclosion, were used for experiments and were used only once. All experiments were performed between ZT = 1-11 at 25°C and 40-60% relative humidity.

### Optogenetic assays – dissection and immunolabeling

Dissections of the male genitalia and female head were performed as described previously with minor modifications ^11^. Briefly, male genitalia were dissected in phosphate-buffered saline (PBS: Takara Bio Inc., #T900; pH 7.4 at 25°C), kept in 50% VECTASHIELD mounting medium (Vector Laboratories, #H-1000; RRID: AB_2336789) in deionized water for ∼5 min, and mounted on glass slides (Matsunami Glass IND., LTD, Osaka, Japan) using VECTASHIELD mounting medium.

Immunolabeling of the brains and ventral nerve cord was performed as described previously with minor modifications ^54^. Briefly, brains were dissected in PBS (pH 7.4 at 25°C), fixed with 4% paraformaldehyde for 60-90 min at 4°C, and subjected to antibody labeling. Brains were kept in 50% glycerol in PBS for ∼1 h, 80% glycerol in deionized water for ∼30 min, and then mounted. Rabbit polyclonal anti-GFP (Invitrogen, #A11122; RRID: AB_221569; 1:1000 dilution) was used for detecting the mCD8::GFP. Mouse anti-Bruchpilot nc82 (Developmental Studies Hybridoma Bank, #nc82, RRID:AB_2314866; 1:20 dilution) was used to visualize neuropils in the brain. Secondary antibodies used in this study were as follows: Alexa Fluor 488-conjugated anti-rabbit IgG (Invitrogen, #A11034; RRID:AB_2576217; 1:300 dilution) and Alexa Fluor 647-conjugated anti-mouse IgG (Invitrogen, #A21236; RRID: AB_2535805; 1:300 dilution).

### Optogenetic assays – confocal microscopy and image processing

Serial optical sections were obtained at 0.84 μm intervals with a resolution of 512 × 512 pixels using an FV1200 laser-scanning confocal microscope (Olympus) equipped with a silicone-oil-immersion 30× lens (UPLSAPO30XSIR, NA = 1.05; Olympus). Images of neurons were registered to the *Drosophila* brain template^55^ by using groupwise registration^56^ with the Computational Morphometry Toolkit (CMTK) registration software. Brain registration, image size, contrast, and brightness were adjusted using Fiji software (version 2.14.0; RRID: SCR_002285).

### Optogenetic assays – retinal feeding

Transgenic male and female flies were maintained under a dark condition for 4-6 days after eclosion and then transferred to a plastic tube (1.5 mL, Eppendorf) containing ∼200 µL of fly food. Plastic tubes that contain males were divided into experimental and control groups. For the experimental group, 2 μL of all-trans-retinal (R2500, Sigma-Aldrich), 25 mg/mL dissolved in 99.5% ethanol (14033-80, KANTO KAGAKU), was placed on the food surface. The male and female flies were kept on the food for 1 day and 2 days, respectively, before being used for the assays.

### Optogenetic assays – a closed-loop system for event-triggered photostimulation

For the event-triggered optogenetic assay (Fig. 5), we used the YORU system on a desktop Windows PC (the same machine used in the speed benchmarking experiments). Green LED light (M530L4, Thorlabs, Inc.) was used as the light source. The cameras used to record the fly behavior and photostimulations, respectively, are as follows: a monochrome CMSO camera (DMK33UX273, The Imaging Source Asia Co., Ltd.) equipped with a light-absorbing and infrared transmitting filter (IR-82, FUJIFILM); a color CMOS camera (DFK 33UP1300, The Imaging Source Asia Co., Ltd). The monochrome CMOS and color CMOS cameras were equipped with a 25 mm focal length lens (TC2514-3MP, KenkoTokina Corporation) and zoom lens (MLM3X-MP, Computar), respectively. The fly behaviors were recorded at a resolution of 640 by 480 pixels and a frame rate of 100 fps for ∼40 min for each fly pair using YORU. The recordings of the photostimulation were performed at a resolution of 1280 × 1080 pixels resolution and 30 fps. The chamber was illuminated from the bottom by an infrared LED light (ISL-150×150-II94-BT, 940 nm, CCS INC.) to enable recordings in dark conditions. We used a round courtship chamber with a sloped wall (19 mm top diameter, 12 bottom diameter, 4 mm height, and 6 mm radius fillet) made of the transparent polylactic acid filament produced by a 3D printer (Sermoon D1, Creality 3D Technology Co., Ltd.). The chamber was enclosed with a slide glass and a white acrylic plate as a lid and bottom, respectively.

For analyzing the behaviors, a male and a female fly were introduced into the chamber by gentle aspiration without anesthesia. YORU captured and detected two behavior object classes. When YORU detected a “wing extension”, a green light (light intensity (530 nm): 2.0 mW/cm^2^) illuminated the entire chamber. As a control experiment of the light condition, we used an event-triggered light with a 1-second delay to illuminate the entire chamber (Fig. 5c). The light intensity of photostimulation was calibrated with an optical power meter (PM100D, Thorlabs, Inc.).

### Optogenetic assays – a closed-loop system for individual-focused photostimulation

For the individual-focused optogenetic assay (Fig. 6), we used the YORU system on a desktop Windows PC (the same as for the speed benchmarking experiments). A projector (HORIZON Pro, XGIMI Technology Co.) was used as a light source to stimulate specific individuals. Camera captures, chambers, and background light setup were configured in the same manner as in the event-triggered photostimulation experiments.

Before the experiments, the positions of the camera and projector planes were calibrated using a circle grid pattern (5 circles height x 8 circles width). The “findHomography” function of “OpenCV” package provided the homography matrix that linearly transformed the position from the camera plane to the projector plane.

For analyzing the behaviors, a male and a female fly were introduced into the chamber by gentle aspiration without anesthesia, and analyzed by YORU. When YORU detected a “wing extension”, a circle-shaped illumination (∼6.5 mm diameter with 1.14 mW/cm^2^ light intensity at 530 nm) was delivered to the individual that was detected as a “fly”. The center coordinates of the predicted bounding boxes were homography transformed using the homography matrix. As a control for the light condition, the projector displayed a patterned light consisting of alternating 3-s of green and 4-s of black”, covering the entire chamber. This was controlled using a custom Python code. This green/black time ratio was determined based on the calculation of courtship duration of wild-type *Drosophila* in courtship assay videos. The light intensity of photostimulation was calibrated with an optical power meter (PM100D, Thorlabs, Inc.).

### Quantification and statistical analysis

Statistical analyses were conducted using Jupyter (Python version 3.9.12) and RStudio (R version 4.3.2). Aligned rank transform one-way analysis of variance (ART one-way ANOVA) was performed to compare the wing extension ratio between conditions in the event-triggered optogenetic assay. All statistical analyses were performed after verifying the equality of variance (Bartlett’s test for three-groups comparisons; F-tests for two-groups comparisons) and normality of the values (Shapiro-Wilk test). Kaplan-Meier curves were generated using R, and a pairwise Log-rank test was performed to compare females’ cumulative copulation rates between conditions in the individual-focused optogenetic assay. After ART one-way ANOVA or pairwise Log-rank tests, p values were adjusted using the Benjamini-Hochberg method in the post hoc test. For the ART one-way ANOVA, the ARTool package (version 0.11.1) was used (https://github.com/mjskay/ARTool/) ^57,58^. For Kaplan-Meier curves and pairwise Log-rank test, the survival package (version 3.5.7) was used (https://github.com/therneau/survival). Statistical significance was set at p < 0.05. Boxplots were drawn using the R package ggplot2 (https://ggplot2.tidyverse.org/). Boxplots represent the median and interquartile range (the distance between the first and third quartiles), and whiskers denote 1.5 × the interquartile range.

## Data and code availability

・ YORU software and all original code have been deposited at GitHub and are publicly available at https://github.com/Kamikouchi-lab/YORU.

## Declaration of generative AI and AI-assisted technologies in the writing process

During the preparation of this work the authors used DeepL (https://www.deepl.com) and ChatGPT (https://chatgpt.com) in order to improve language. After using this tool, the authors reviewed and edited the content as needed and take full responsibility for the content of the publication.

## Acknowledgments

We thank Mayu Yamanouchi for the software logo design; Masahiko Hibi and Shiori Hosaka for collecting animal behavior videos; Ryota Nishimura (the Research Equipment Development Group, Technical Center of Nagoya University) for the chambers used in behavioral experiments, and construction virtual reality environment for mice; Matthew P. Su, Kentaro Noma, Tomoka Nozaki and Haruka Ando for discussions; Kei Ito, Daniel F. Eberl and the Bloomington Drosophila Stock Center (Indiana University, Bloomington, IN, USA) for fly stocks; Developmental Studies Hybridoma Bank for antibodies. This study was financially supported by the MEXT KAKENHI Grant-in-Aid for Transformative Research Areas (A) “iPlasticity” (JP23H04228 to A.K.), Grant-in-Aid for Transformative Research Areas (A) Hierarchical Bio-Navigation (JP22H05650 and JP24H01433 to R.T.), Grants-in-Aid for Scientific Research (C) (JP23K05846 to R.T. and JP23K05845 to T.S.), Grant-in-Aid for Early-Career Scientists (JP20K16464 to R.F.T. and JP21K15137 to R.T.) JST FOREST (JPMJFR2147 to A.K.), the Japan Society for the Promotion of Science (JSPS) Grant-in-Aid for JSPS Fellows (No. JP24KJ1290 to H.M.Y.) Japan.

## Supplemental file legends

**Supplementary Table 1: Summary of model accuracy compared to human annotations**

**Supplementary Table 2: Summary of model Precision and Recall in each pretrained model and in each training image number**

**Supplementary Table 3: Summary of model accuracy of “mouse – treadmill” model**

**Supplementary Table 4: Summary of inference speed of one frame detection**

**Supplementary Table 5: Summary of dataset conditions**

**Supplementary Table 6: Lists of PC hardware information for benchmarking tests**

