## Supplementary_figures.pdf for "YORU: social behavior detection based on user-defined animal appearance using deep learning"

Figure S1

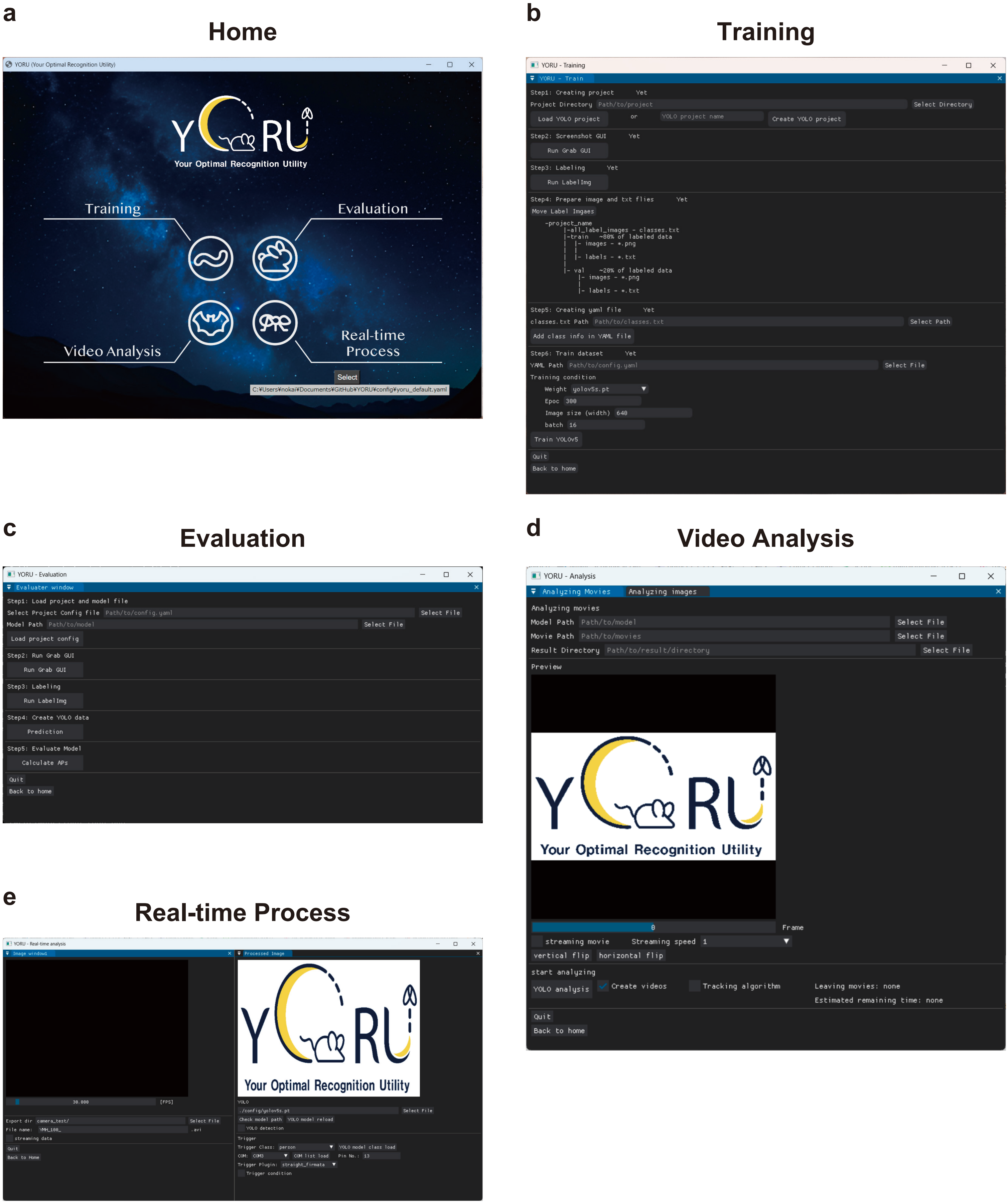


**Figs. 1: YORU's GUI screenshots**
**a-e**, GUI screenshots of "Home" (**a**), "Training" (**b**), "Evaluation" (**c**), "Video Analysis" (**d**), and "Real-time Process" (**e**) packages.

Figure S2

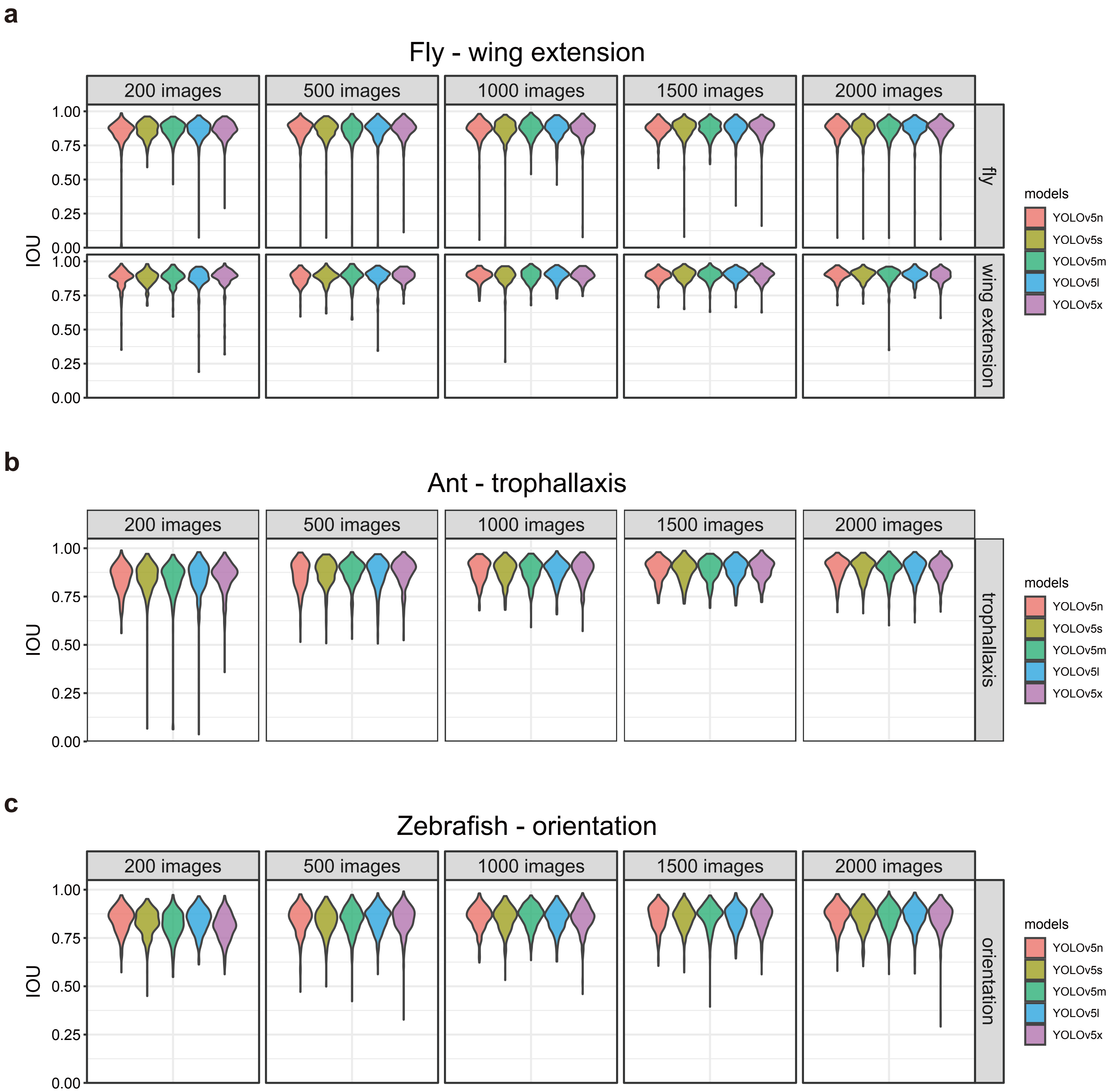


**Figs. 2: IOU distributions of three individual social behaviors.**
**a-c**, IOU distributions of “Fly - wing extension” (**a**), “Ant - trophallaxis” (**b**), and “Zebrafish - orientation” (**c**) models based on each YOLOv5 pre-trained model. Violin plots represent the probability density of individual data points within the range of possible values (Also in the following Figures).

Figure S3

**a**

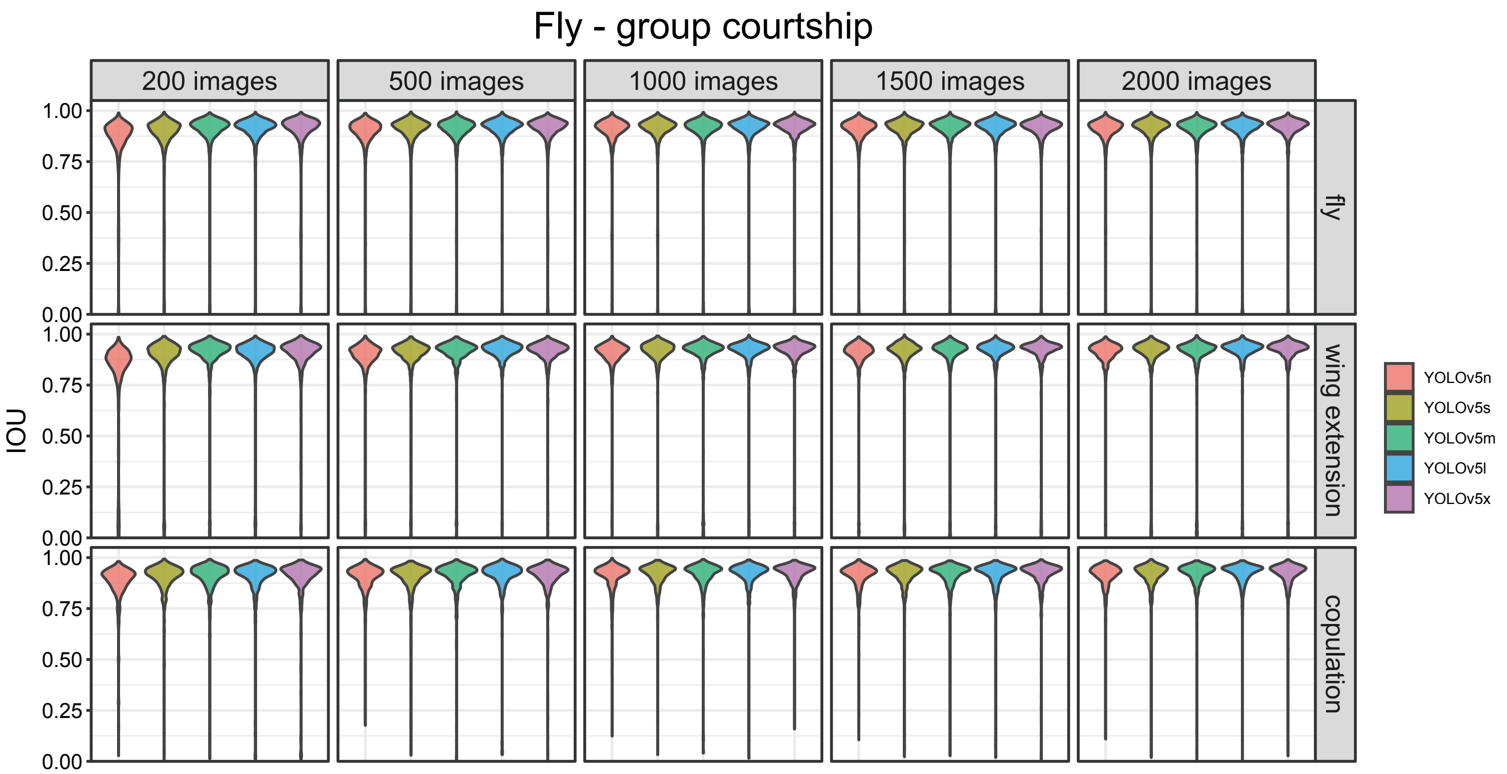


**b**

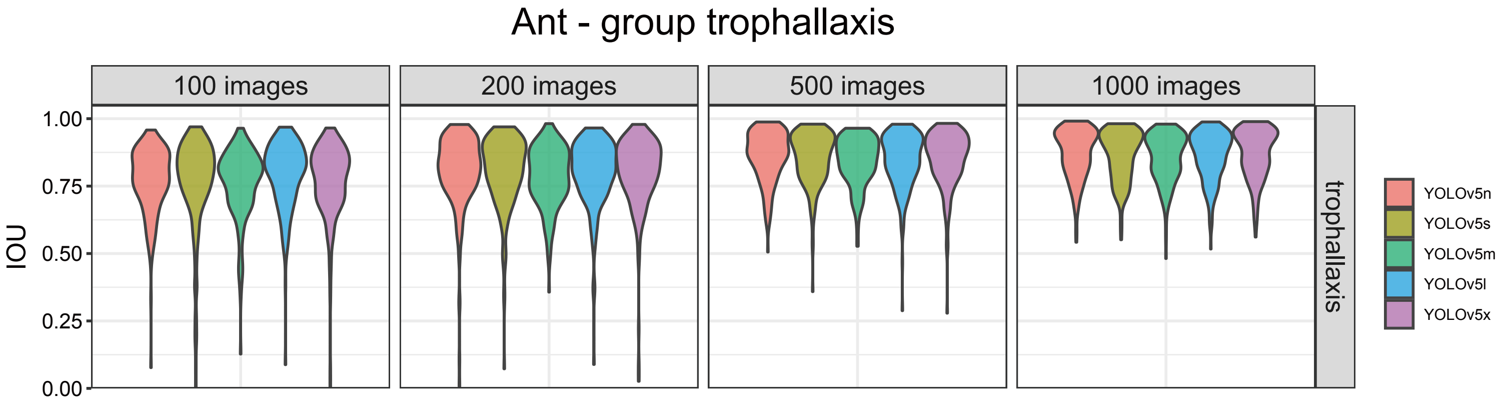


**Figs. 3: IOU distributions in multi-individual conditions.**
**a**,**b**, IOU distributions of “Fly - group courtship” (**a**) and “Ant - group trophallaxis” (**b**) models based on each YOLOv5 pre-trained model.

Figure S4

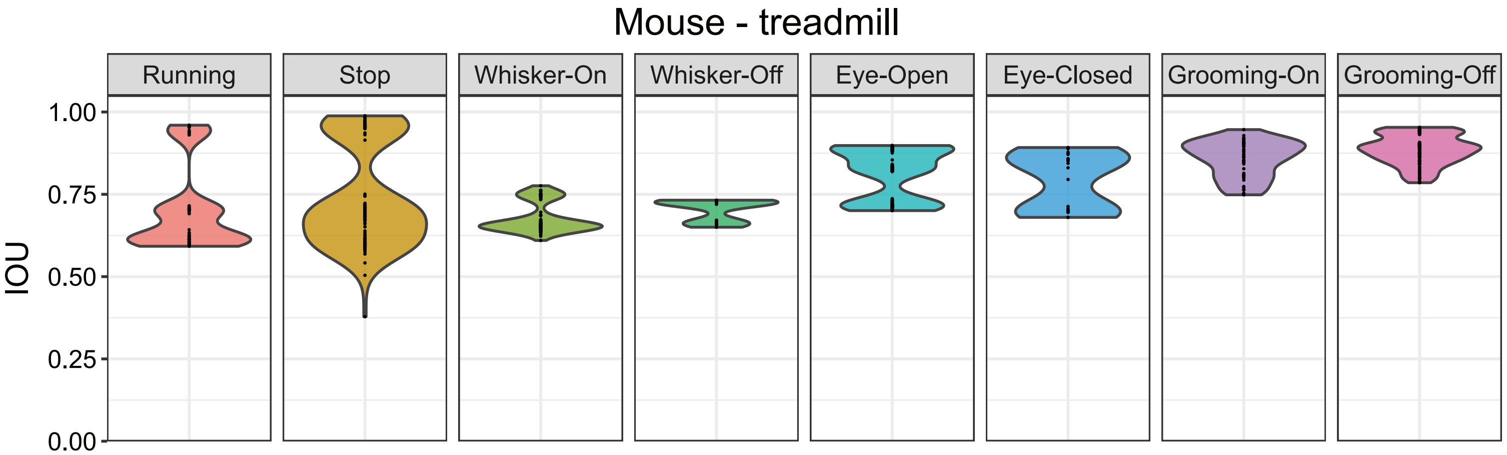


**Figs. 4: IOU distributions of mouse behaviors.**
IOU distributions of “Mouse - treadmill” model based on YOLOv5s pre-trained model.

Figure S5

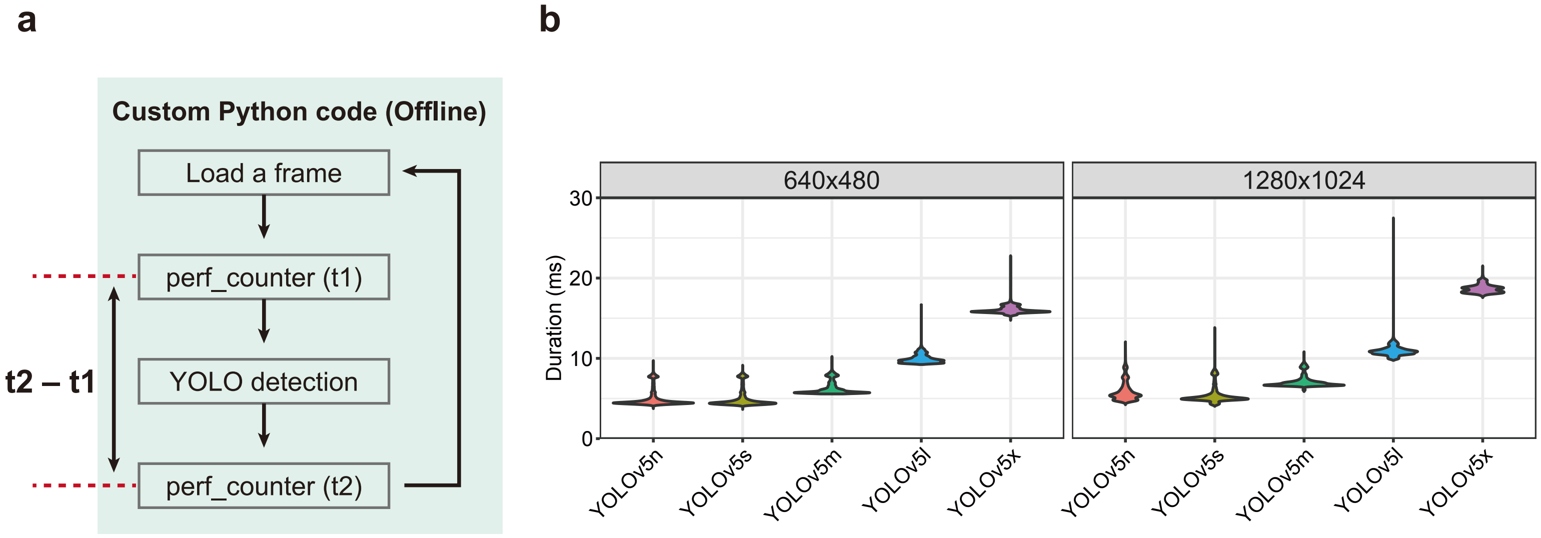


**Figs. 5: YORU’ s inference speed of different networks and image sizes.**
**a**, Schematic of inference speed calculation of YORU’ s object detection. In the custom Python code in YORU’ s “Real-time Process” package, the times before and after the YOLO detection step (t1 and t2, respectively) were logged. The inference speed of one frame detection was calculated as the difference between t1 and t2.
**b**, Inference speeds of “LED lighting - Small” (Left) and “LED lighting - Large” (Right) models for frame detection. The system latency of each model was calculated from camera images with resolutions of 640x480 pixels and 1240x1024 pixels, respectively.

Figure S6

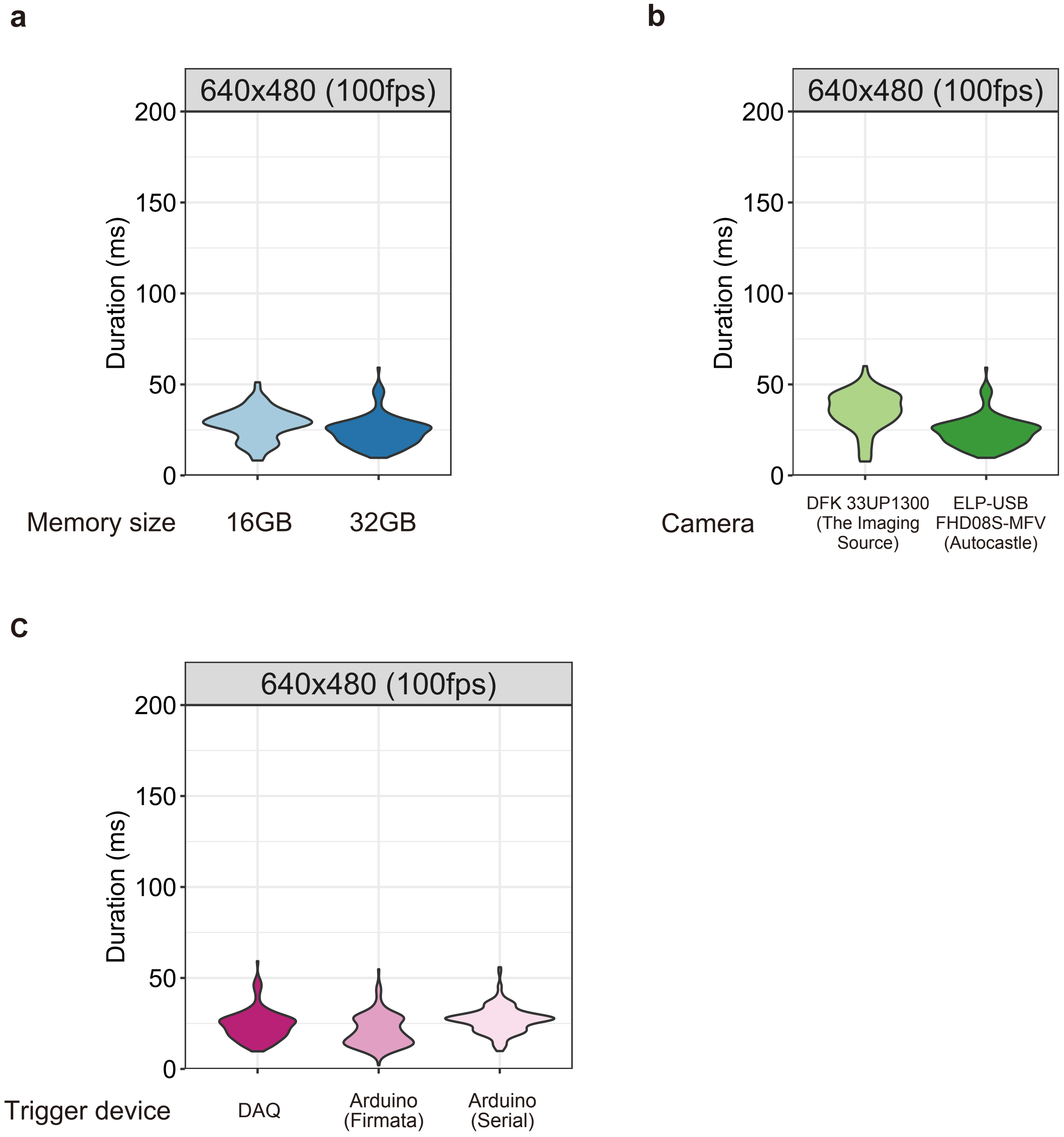


**Figs. 6: System latency in different system’ s hardware.**
**a-c**, System latency of different PC memory sizes (**a**), cameras (**b**), and trigger types (**c**).

Figure S7

*JO15-2-GAL4 >*
*20xUAS-IVS-mCD8::GFP*
(Female)

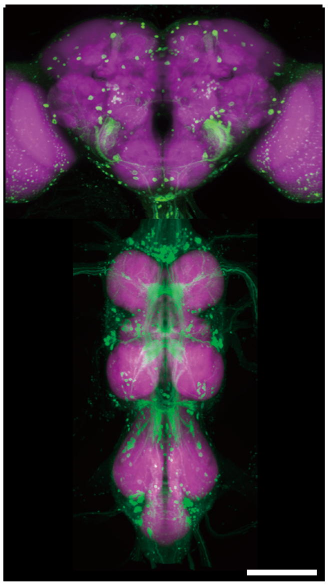

**Figs. 7: Expression pattern of JO15-2-GAL4 in a female fly.**
*JO15-2-GAL4* expression in the female brain and ventral nerve cord. GFP markers driven by *JO15-2-GAL4* (*JO15-2-GAL4>20XUAS-IVS-mCD8::GFP*) are detected in the brain and ventral nerve cord. Scale bar, 100μm. Signals of the GFP marker (green) and counter-labeling with the nc82 antibody (magenta) are shown.
